# Identification of dehydrin protein complexes *in vivo* reveals functional interactions of LEA5 with OSCA3, PIP2B and PLD*α*1 in plant water-deficit stress

**DOI:** 10.1101/2025.10.27.684818

**Authors:** Norma Fàbregas, Félix Juan Martinez Rivas, Itzell Eurídice Hernández-Sánchez, Fidel Lozano-Elena, Fayezeh Arabi, Ewelina Sokolowska, Aleksandra Skirycz, Alisdair R. Fernie

**Author notes:** These authors contributed equally: Norma Fàbregas, Félix Juan Martinez Rivas.

## Abstract

Climate change and especially the concomitant increasing frequency of drought and salinity induced water-deficit stresses, is a major limiting factor of plant growth and crop productivity worldwide. Understanding the adaptive responses of plants to water-deficit stress has therefore become a central challenge in plant biotechnology. Considerable evidence indicates that late embryogenesis abundant (LEA) proteins contribute to water-deficit stress tolerance and the stabilization of metabolic enzymes, yet the molecular basis of these responses remains unknown. To date, limited direct evidence for specific *in vivo* interactions between LEA proteins and their molecular targets has been reported. Here, we identify for the first time the native *in vivo* interactome of three Arabidopsis dehydrin LEA proteins - LEA4 (COR47), LEA5 (ERD10), and LEA10 (ERD14) - expressed under their own promoters and in response to salt stress. Our results show that LEA4, LEA5, and LEA10 dehydrins interact with each other, and share six common candidate protein interactors, suggesting that they form a core protein complex in salt stress conditions. We confirmed the direct protein-protein interaction between LEA5 with LEA4, LEA10, OSCA3, PLD*α*1, PIP2B and OST1/AHA1 at the biochemical level. Phenotypic analyses of loss-of-function genetic mutants revealed that the interactions of LEA5 with OSCA3 and PIP2B promote seed germination under salt stress. LEA5 interaction with PLDα1 promotes root growth under salt stress, while its interaction with PIP2B enhances root growth under osmotic stress, indicating distinct stress-specific functional roles for each interaction. In conclusion, this study identifies the native *in vivo* interactions of three dehydrin proteins, while uncovering the functional relevance of these interactions under salt and osmotic stress conditions, thus providing novel mechanistic insights into the role of LEA proteins in water-deficit stress adaptation.

## INTRODUCTION

Climate change is one of the most urgent global challenges, and involves more frequent and intense droughts, as well as high salinity levels in agricultural soils. These water-deficit environmental stressors are significant threats to ecosystems and are detrimental for agricultural productivity (Gupta et al. 2020). Drought, characterized by prolonged periods of water scarcity, and salinity, defined by an excessive accumulation of soluble salts in soil, both disrupt seed germination and plant growth, diminishing crop yields, and threaten food security (Lesk et al. 2016; Gupta et al. 2020; Van Zelm et al. 2020). Addressing the challenges posed by water-deficit stresses such as drought and salinity is thus critical to ensuring sustainable agricultural systems in the face of climate change. It is also of outmost importance to breed crops capable of tolerating these water-deficit stresses without a significant impact on their yield. However, our knowledge concerning drought and salinity stress resistance is still limited, partly due to the complexity of the trait, involving important metabolic and physiological changes at the whole plant level. Therefore, it is of paramount importance to elucidate the mechanisms by which plants respond to and adapt to drought and salt stress.

High salinity and drought stress have synergistic effects on plant physiological processes, leading to reduced water uptake, impaired nutrient absorption, and disrupted metabolic activities (Munns and Tester 2008). Both drought and salinity cause osmotic stress by lowering the water potential of plant cells, which can lead to cell turgor loss, membrane disorganization, protein denaturation, inhibition of photosynthesis and oxidative damage, and thus both stresses share crosstalk physiological and molecular plant responses (Ma et al. 2020). Osmotic stress promotes the accumulation of abscisic acid (ABA) in plant cells and the activation of the ABA signalling pathway, but also the activation of an ABA-independent pathway. To date, many signal transduction components of the ABA-dependent (Park et al. 2009; Santiago et al. 2009; Cutler et al. 2010) and the ABA-independent pathways (Seki et al. 2007; Yoshida et al. 2010; Hauser et al. 2011) have been identified. Recent studies have identified osmotic-stress-activated Raf-like protein kinases as upstream regulators of ABA-dependent and -independent signalling pathways in *Physcomitrella patens* (Saruhashi et al. 2015) and Arabidopsis (*Arabidopsis thaliana*) (Katsuta et al. 2020; Lin et al. 2020; Soma et al. 2020; Takahashi et al. 2020). These studies have brought new insights into the molecular understanding of water-deficit stress tolerance in both Arabidopsis and crop species (Fàbregas et al. 2020).

Although many genes, proteins and metabolites connected to water-deficit stress adaptation responses have been identified, the functional relevance for most of these molecules is poorly understood. A clear example of this is the case of the LATE EMBRYOGENESIS ABUNDANT (LEA) proteins, which accumulate in response to water-deficit stress responses. However, the molecular and functional relevance of the stress-accumulation of LEA proteins *in vivo* remains an open question.

LEA proteins were first discovered in cotton seeds (Galau et al. 1986) and later in other higher plants and mosses, bacteria, nematodes, fungi and animals (Hand et al. 2011). LEA proteins are highly hydrophilic and belong to a large family of intrinsically disordered proteins (IDPs), which, upon desiccation, become structurally stabilized (Mouillon et al. 2006). IDPs show high specificity and low binding affinity protein-protein interactions during signal transduction (Uversky et al. 2005), facilitating the formation of transient and flexible interaction networks (Oldfield et al. 2008). Moreover, IDPs are critical for plant adaptation in challenging environments because of their ability to perform multiple functions, such as antiaggregating, protein and membrane stabilization, as well as molecular chaperone-like activities (Chakrabortee et al. 2007; Battaglia et al. 2008; Kovacs et al. 2008; Hincha and Thalhammer 2012; Cuevas-Velazquez et al. 2017).

LEA proteins protect other proteins from aggregation caused by desiccation and osmotic water-deficit stresses in animals and plants (Goyal et al. 2005). In plants, LEA proteins show different expression patterns throughout plant tissues participating in different regulatory pathways (Bies-Ethève et al. 2008). The underrepresentation of LEA genes in the desiccation-sensitive marine water species *Zostera marina* and freshwater species *Spirodela polyrhiza* (Olsen et al. 2016), and the overrepresentation of LEA genes in the desiccation-tolerant species *Xerophyta viscosa* (Costa et al. 2017) suggest that the evolution of LEA genes contributed to water-deficit stress adaptation in plants. However, although various experimental observations imply that LEA proteins participate in the response and adaptation to water-deficit stress (Tunnacliffe et al. 2010; Hand et al. 2011; Hernández-Sánchez et al. 2022), the molecular basis and biochemical functions of most LEA proteins remain largely unknown.

In Arabidopsis, 51 LEA proteins have been identified, divided into eight heterogeneous families (Hundertmark and Hincha 2008). In literature, different nomenclatures have been used to refer to the LEA proteins. We have summarized the more commonly used nomenclatures, the family groups and the subcellular localization of LEA proteins in Supplementary Table 1. In this work, for the sake of clarity, we used the nomenclature of LEA genes that have been systematically named according to their positions within the genome, sequentially ranging from LEA1 to LEA51 (Hundertmark and Hincha 2008). The largest family of LEA proteins is LEA_4 with 18 members, followed by the dehydrin group, with 10 members, the SMP group, with six members, with the rest of family groups (LEA_1, LEA_2, LEA_3, LEA_5, and LEA_6) being represented by four or less members (Supplementary Table 1). LEA_5 and SMP families appeared early in the plant lineage, while other families emerged later in plant evolution (Artur et al. 2019). Transient expression analyses of the 51 LEA proteins in Arabidopsis protoplasts revealed that most LEA proteins are localized to the cytosol (70%) while the rest are variously localized in chloroplasts and mitochondria, vacuoles, endoplasmic reticulum and pexophagosomes (Candat et al. 2014) (Supplementary Table 1).

Dehydrins are a well characterized group of LEA proteins (Graether 2022). All dehydrin proteins are characterized to have a common lysine-rich domain (K-segment), which is typically located in the C-terminal position, and specific antibodies against this domain have been classically used to identify related proteins (Close 1997). Moreover, a serine stretch (S-segment) is present in eight members of the dehydrin family (Close 1996). The S-segment is phosphorylated *in vivo* (Goday et al. 1994) and can bind calcium (Alsheikh et al. 2005) whereas the K-segment is essential for protective functions (Yang et al. 2015). Two N-terminal conserved consensus sequences have been identified in the dehydrin family. The Y-segment is a six-residue motif (DEYGNP) present in three dehydrin members in Arabidopsis (Close 1996; Malik et al. 2017). The more recently identified F-segment (DRGLFDFLGKK) is present in four dehydrin proteins lacking the Y-segment in Arabidopsis (Richard Strimbeck 2017). A less common motif H-segment is present in one single dehydrin (Melgar and Zelada 2021). Phosphoproteomics studies identified a group of LEA proteins that are phosphorylated in Arabidopsis seeds (Irar et al. 2006). Three dehydrin proteins, LEA4 (COR47), LEA5 (ERD10), and LEA10 (ERD14) in Arabidopsis and the dehydrin RAB17 in maize are phosphorylated *in vivo* by the Casein Kinase 2 (Goday et al. 1994; Alsheikh et al. 2003; Riera et al. 2004). Interestingly, phosphorylation regulates binding of dehydrin proteins to calcium ions *in vivo* (Alsheikh et al. 2003, 2005), as well as membrane binding (Eriksson et al. 2011) and subcellular localization (Riera et al. 2004). Dehydrin proteins accumulate in response to different abiotic stress conditions, including dehydration, heat and cold stress, osmotic stress, salt stress, ABA treatment and drought imposed by withholding water (Gilmour et al. 1992; Nylander et al. 2001; Riyazuddin et al. 2022; Szlachtowska and Rurek 2023).

LEA proteins bind to a wide variety of macromolecules ranging from nucleoproteins and membrane complexes to sugars (Close 1997). Protective chaperone activity has been shown for LEA5 (ERD10) and LEA10 (ERD14) dehydrin proteins (Kovacs et al. 2008). Moreover, other LEA proteins stabilize key metabolic enzymes upon dehydration stress in different plant species (Hara et al. 2001; Bravo et al. 2003; Grelet et al. 2005; Reyes et al. 2005). Within the LEA_2 family, LEA1 and LEA27 protect lactate dehydrogenase from inactivation during desiccation-rehydration (Dang et al. 2014; Popova et al. 2015). In addition, LEA proteins can stabilize fumarase and citrate synthase enzymes from wheat and nematodes *in vitro* (Goyal et al. 2005; Grelet et al. 2005). Altogether, these studies demonstrate that LEA proteins can stabilize labile enzymes under stress conditions *in vitro*.

Based on the phenotypic analyses of loss-of-function genetic mutants, a few LEA proteins have been implicated in growth and survival under osmotic stress conditions. The *knock-out* mutants for the dehydrin LEA33 (Lti30) are hypersensitive to ABA and overexpression lines are resistant to ABA (Shi et al. 2015). In agreement, LEA33 (Lti30) overexpression lines are tolerant to cold and freezing and accumulate proline (Puhakainen et al. 2004). Moreover, LEA5 (ERD10) dehydrin is induced by cold treatment and *erd10* ko mutants are sensitive to cold stress and drought stress assays in soil (Kim and Nam 2010). However, despite growing evidence of the physiological roles of LEA proteins, the molecular interactors and mechanisms underlying their function in plants remain largely unknown. This is partly due to the intrinsically disordered nature of LEA proteins, which lack stable tertiary structures under physiological conditions, making them difficult to study using conventional structural biology approaches. Additionally, their interactions with target molecules are often transient, condition-dependent, and of low affinity, posing further challenges for *in vivo* detection and characterization. These properties, while likely essential for their protective functions under stress, contribute to the complexity of elucidating their precise roles at the molecular level.

In this context, we hypothesize that certain LEA proteins from the dehydrin family function as molecular hubs that coordinate signalling and metabolic pathways required for plant stress adaptation. We focus on dehydrins due to their capacity for post-translational modification, such as phosphorylation, and their protective enzymatic activities, which make them strong candidates for linking stress signaling with metabolic regulation. Here, we disclose *in vivo* protein-protein interactions of three LEA proteins from the dehydrin family with other stress-related proteins in the model plant Arabidopsis, and we validate some of these interactions through genetic approaches. This work provides new insights into the molecular mechanisms by which dehydrin LEA proteins contribute to plant responses under water-deficit conditions, including salt and osmotic stress.

## RESULTS

### Dehydrins LEA4, LEA5 and LEA 10 accumulate in response to salt stress

In the present work, we focus on the study of the dehydrin family in Arabidopsis that includes ten members. According to publicly available microarray data (Schmid et al. 2005), half of the dehydrin genes are predominantly expressed in seeds, while the other half show high expression in vegetative tissues such as roots and stems (Table 1). All ten dehydrin proteins are localized in the cytosol (Candat et al. 2014) and four of them can be phosphorylated (Alsheikh et al. 2005). Publicly available transcriptomic data further reveal the response of dehydrin genes to specific water-deficit stress treatments: LEA5, LEA10, LEA45 and LEA51 are induced by drought; LEA4, LEA5, LEA10 and LEA33 are induced by ABA; and LEA4, LEA5, LEA10, LEA33, LEA34, LEA44, LEA45 and LEA51 are induced by salt stress (Table 1). We selected LEA4, LEA5, and LEA10 for further analysis because they are highly expressed in vegetative tissues, are phosphorylated, and show consistent induction under different water-deficit stress (as highlighted in Table 1), making them strong candidates for investigating our working hypothesis.

**Table 1.**
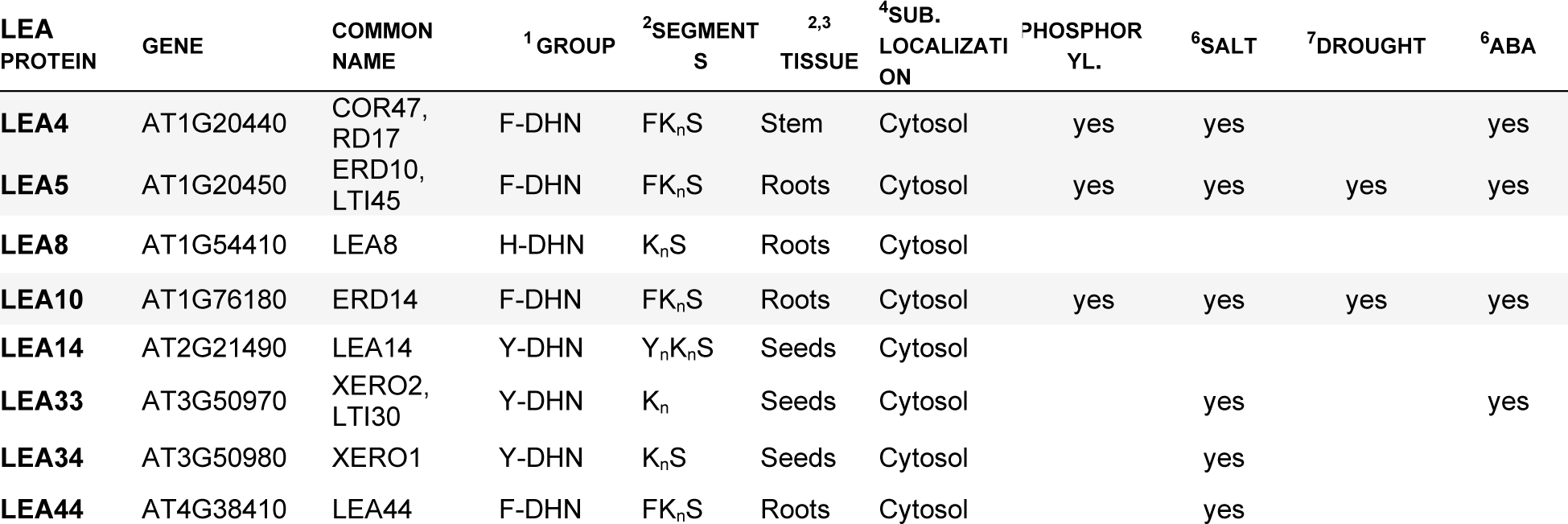
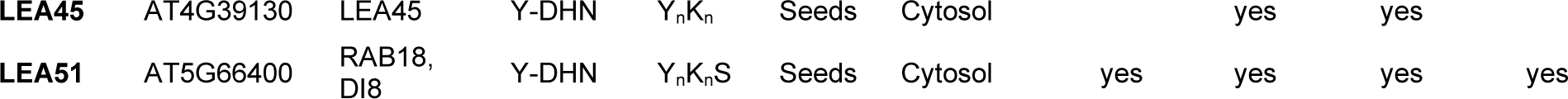
Dehydrin proteins family localization and transcriptional response to stress. Data extracted from **^1^**(Melgar and Zelada 2021); **^2^**(Hundertmark and Hincha 2008); **^3^**(Schmid et al. 2005); **^4^**(Candat et al. 2014); **^5^**(Alsheikh et al. 2005); **^6^**(Kilian et al. 2007); **^7^**(Pandey et al. 2010)

Furthermore, we analysed further public microarray data (Schmid et al. 2005; Brady et al. 2007), which revealed similar expression patterns of mRNAs for LEA4, LEA5 and LEA10 in different developmental stages, and particularly in the primary root, where they exhibited enriched expression in the cortex and vascular tissues in control conditions (Supplementary Figure 1). LEA4, LEA5 and LEA10 transcripts displayed a similar spatiotemporal response to salt stress (Dinneny et al. 2008) and ABA treatments (Pandey et al. 2010) (Supplementary Figure 2). The LEA4, LEA5 and LEA10 transcripts accumulated in the cortex and vascular tissues of the root under salt stress and in the stomatal guard cells following ABA treatment (Supplementary Figure 2). In contrast, other members of the dehydrin protein family, such as LEA44 and LEA51, showed either very low or no response to salt stress and a more restricted response to ABA (Supplementary Figure 3).

In summary, we primarily focused on the characterization of three dehydrin family members - LEA4, LEA5 and LEA10 – because previous available data indicate that (i) they are not seed specific *i.e.* they are higher expressed and in a broader type of cells and tissues than other dehydrins in control conditions, (ii) they can be phosphorylated, (iii) they showed a similar response to salt stress in terms of mRNA levels.

To investigate the molecular function of dehydrin proteins, we generated Arabidopsis translational reporter lines with the Green fluorescent protein (GFP fused to LEA proteins and expressed under the control of their native promoters (*pLEA:LEA-GFP*, Material and methods). In agreement with previous transcriptomic data, LEA4, LEA5 and LEA10 showed basal protein levels in Arabidopsis seedlings at control conditions (Figures 1 and 2). Importantly, a similar and significant accumulation of LEA4, LEA5 and LEA10 proteins was observed after 1 hour of salt stress treatment, which continued to increase within the time course until the last timepoint at 24 hours of salt treatment (Figure 1). In contrast, other dehydrin proteins family members, such as LEA44 and LEA51, showed a lack of response to salt stress although they were accumulated upon ABA treatment (Supplementary Figure 4), in agreement with previously reported mRNA levels (Supplementary Figure 3). The stress-dependent accumulation of LEA4, LEA5 and LEA10 dehydrin proteins expressed under their respective native promoters was most prominent at 24 hours of salt stress in comparison to ABA treatment (Figure 1). Thus, in this study, we focused on the characterization of functions and interactions of LEA4, LEA5 and LEA10 dehydrin proteins under salt stress conditions.

**Figure 1.**
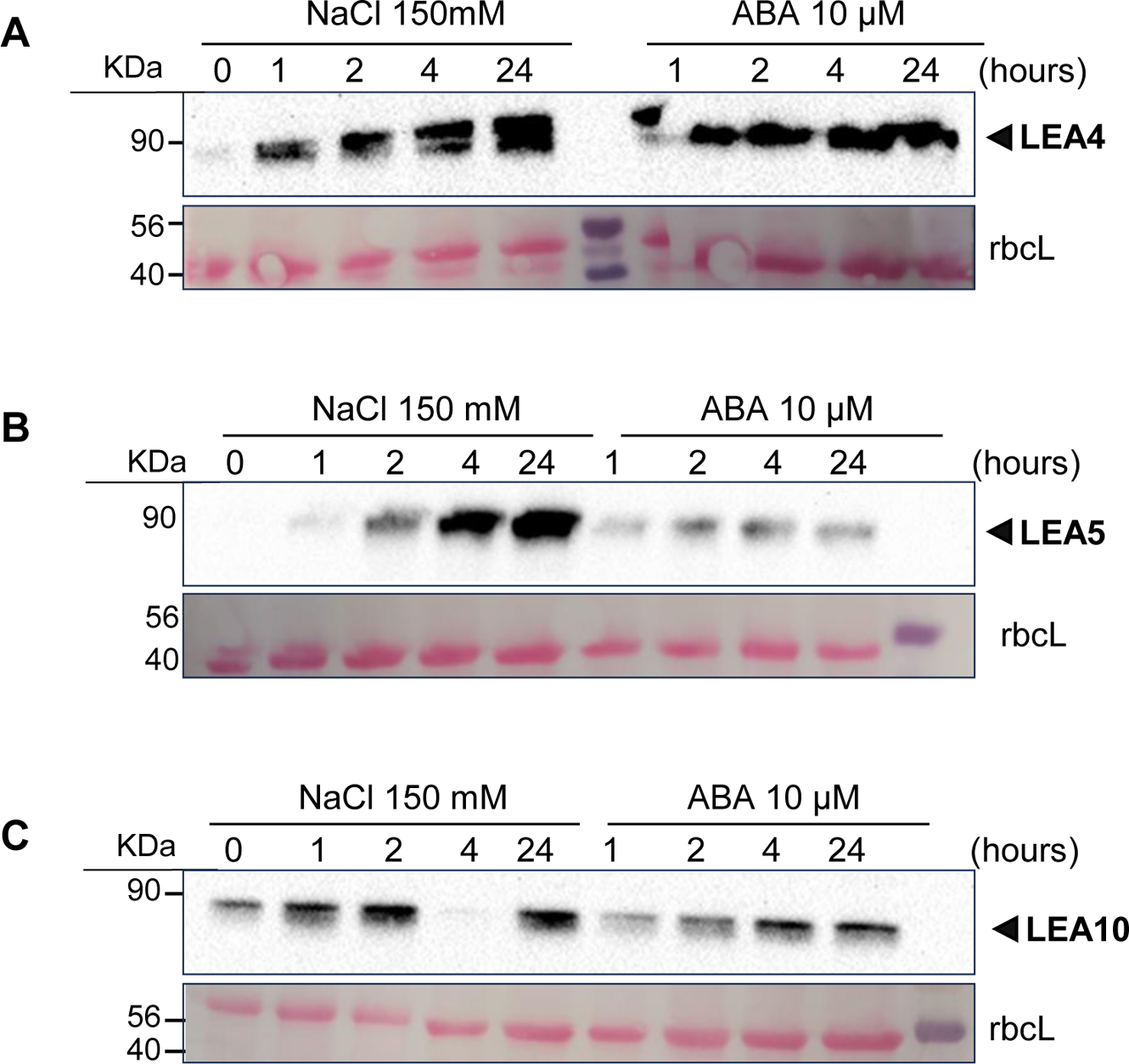
Protein levels of LEA4, LEA5, and LEA10 dehydrins in response to NaCl and ABA treatments. Immunoblot of total protein extracts from (A) *pLEA4:LEA4-GFP*, (B) *pLEA5:LEA5-GFP* and (C) *pLEA10:LEA10-GFP* 10-day-old Arabidopsis seedlings in response to salt stress (150 mM NaCl) or ABA treatment (10 µM). Samples were collected at different time points of the treatment as indicated in hours. For the western blot, anti-GFP antibodies were used in total protein extracts of T3 homozygous translational lines. Top row: immunoblot with antiGFP primary antibody. Bottom row: Ponceau S staining of the Rubisco large subunit (rbcL) as a control of protein loading and transfer. LEA4 molecular weight is 29,9 KDa. LEA5 molecular weight is 29,5 KDa, LEA10 molecular weight is 20,79 KDa and GFP molecular weight is 25 KDa.

**Figure 2.**
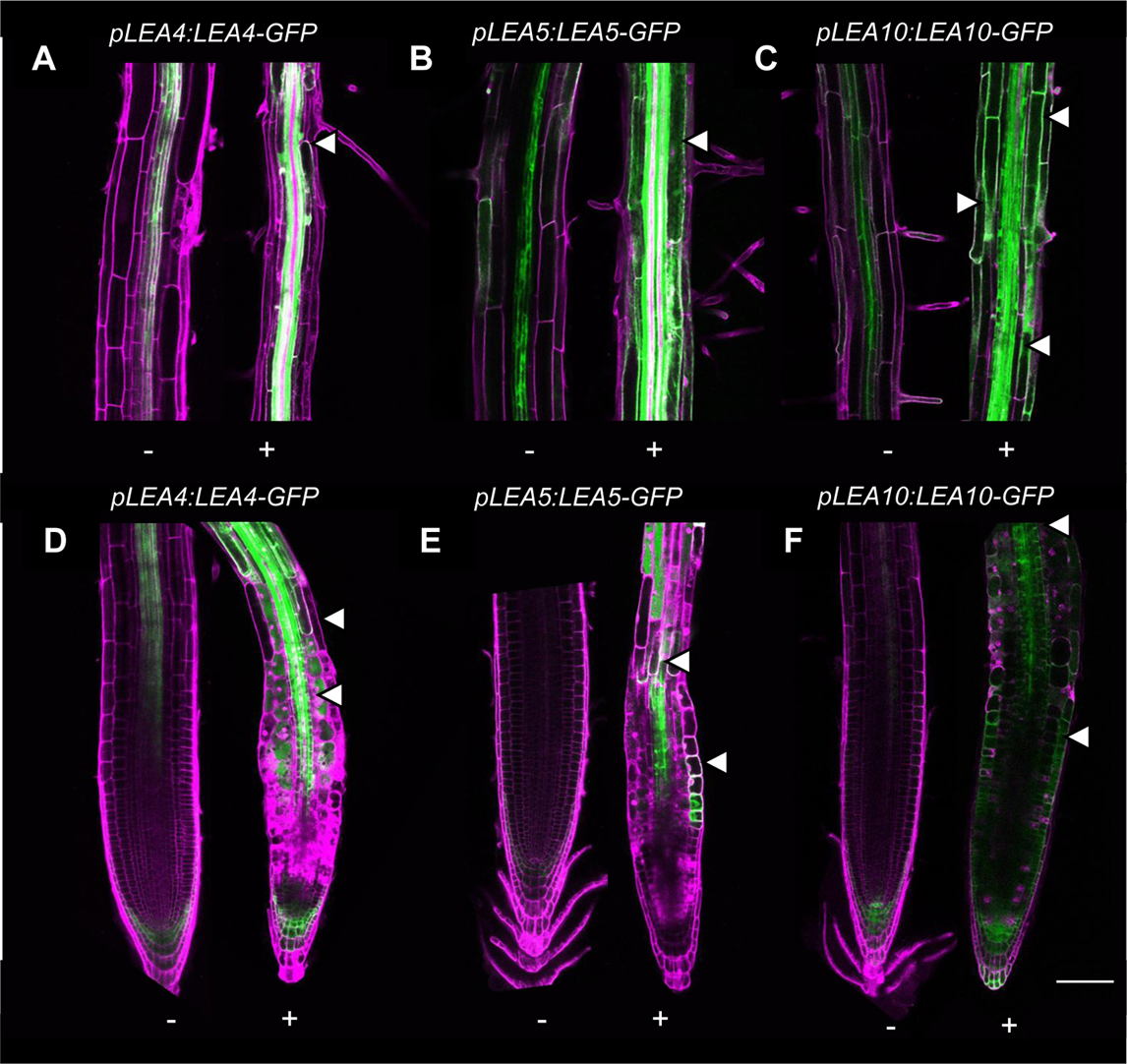
LEA4, LEA5 and LEA10 proteins accumulate under salt stress and expand their spatial localization in the primary root. Confocal imaging of roots of (A, D) *pLEA4:LEA4-GFP*, (B, E) *pLEA5:LEA5-GFP* and (C, F) *pLEA10:LEA10-GFP* 10-day-old seedlings. Upper panel shows the root differentiation zone (A, B, C) and lower panel shows the root tip (D, E, F). Green channel depicts the protein localization of LEA4, LEA5, and LEA10 at the vascular tissues of the root in control conditions (-) and under 24 h of salt stress 150 mM NaCl (+), expands to other cell types and tissues as indicated by white arrows. Magenta channel depicts the propidium iodine staining of the plant cell wall, which also stains the nuclei of death cells under stress (+) as indicated by white arrows. Scale bar: 100 µM.

Since LEA proteins are naturally disordered in control conditions, setting up appropriate stress conditions is crucial to increase LEA protein levels, and stabilize LEA protein structures and their transient and low affinity interactions. For these reasons, characterization of LEA4, LEA5 and LEA10 protein complexes was performed in plants exposed to salt stress (150 mM NaCl) for 24 hours.

*In vivo* protein localization analyses using confocal microscopy in Arabidopsis roots, revealed the preferential accumulation of LEA4, LEA5 and LEA10 proteins in the vascular tissues of the primary root (Figure 2). While LEA4 and LEA10 specifically localized to the root vascular tissues, LEA5 showed high accumulation in the vasculature but was also present at lower levels in the epidermis and cortex cells of the root elongation zone in control conditions (Figure 2). Upon 24h of salt stress, LEA4, LEA5 and LEA10 proteins levels increased in the vascular tissues of roots meristem and elongation zones, and their localization pattern expanded to other root cell types, such as the epidermis and cortex layers (Figure 2).

### Dehydrin proteins LEA4, LEA5 and LEA10 form a core protein complex in salt stress conditions

To identify the interacting proteins of the LEA4, LEA5 and LEA10 dehydrins in Arabidopsis, we performed *in vivo* immunoprecipitation of LEA4, LEA5 and LEA10 protein complexes in plants exposed to salt stress (150 mM NaCl) for 24 hours, ensuring elevated levels of the stressed-dehydrin bait proteins. Protein extracts were immunoprecipitated with anti-GFP antibodies from the corresponding *pLEA:LEA-GFP* lines. The *pLEA10::GFP* line was included as a negative control. The immunoprecipitates were subsequently analysed using liquid chromatography–tandem mass spectrometry (LC-MS/MS, Material and Methods). A total of 160 LEA4-associated proteins in *pLEA4:LEA4-GFP* lines (Supplementary Table 2), 331 LEA5-associated proteins in *pLEA5:LEA5-GFP* lines (Supplementary Table 3) and 93 LEA10-associated proteins in *pLEA10:LEA10-GFP* lines (Supplementary Table 4), were identified in at least two out of the three immunoprecipitation biological replicates. Only proteins represented by at least two unique peptides were considered. The LEA4_1 sample showed only five protein candidates, in contrast to the 160 identified in the other two replicates, indicating it as a clear outlier (Supplementary Table 2). Consequently, the LEA4_1 sample was excluded from further analyses. Proteins detected in the negative control line *pLEA10::GFP* were subtracted by calculating the ratio of protein abundance between the samples and the negative control, and by filtering the candidate proteins with a fold change greater than 1.5. By this means, 16 candidate interacting proteins of LEA4 (Supplementary Table 5), 81 candidate interacting proteins of LEA5 (Supplementary Table 6), and 19 candidate interacting proteins of LEA10 (Supplementary Table 7) were identified as high-confidence candidate interactors excluding the GFP tag protein.

Six common candidate protein interactors were identified in all three LEA4, LEA5 and LEA10 protein complexes (Figure 3A,B). Among these candidates, LEA10 dehydrin protein together with other five water-deficit stress-related proteins were identified: Two aquaporin PLASMA MEMBRANE INTRINSIC PROTEINS (PIP2A, PIP2B), a proton pump ATPase (AHA1, also known as OST2), a REMORIN family protein (REM1.3), and a PLASMA MEMBRANE ASSOCIATED CATION-BINDING PROTEIN 1 (PCAP1). Gene Ontology (GO) enrichment analyses confirmed that the six common detected proteins in LEA4, LEA5 and LEA10 immunoprecipitations were involved in responses to water-deficit stress (Figure 3C,D). Remarkably, GO enrichment analysis performed on the individual LEA protein complexes revealed that distinct biological functions are associated with each complex, suggesting functional specialization among the LEA proteins. Associated proteins to LEA4, LEA5, and LEA10 were enriched in microtubule/cytoskeleton organization, metabolic processes, and stomatal closure functions, respectively, among others (Figure 3E). However, cold and water transport GO categories were highly enriched in the three protein complexes, suggesting that LEA interactor proteins may have evolved specialized functions while preserving a water-deficit stress response core.

**Figure 3.**
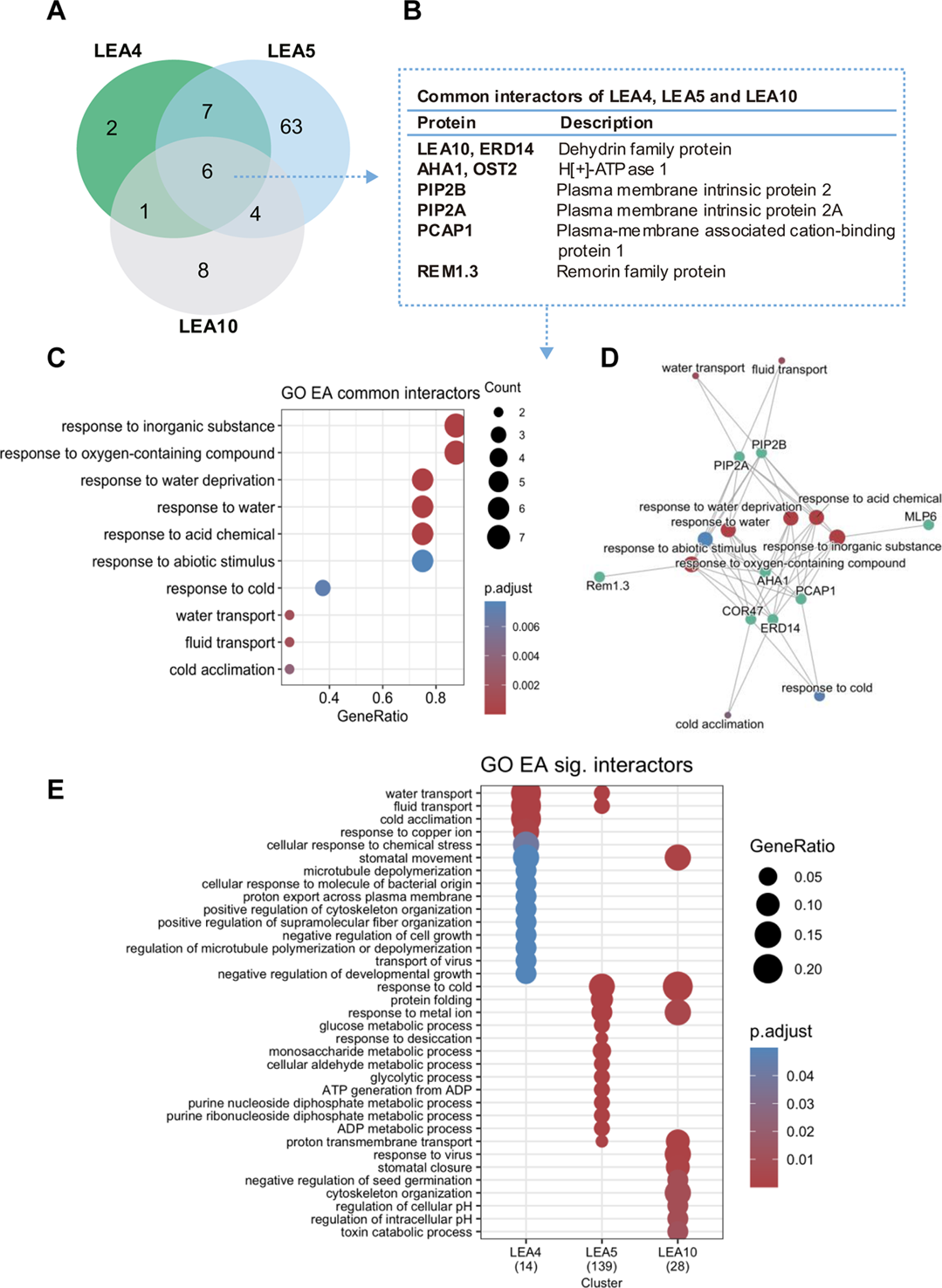
Gene Ontology Enrichment Analysis of LEA4, LEA5 and LEA10 protein interactors under salt stress. (A) Venn Diagram comparing lists of candidate interactors: 16 in LEA4 immunoprecipitates, 81 in LEA5 immunoprecipitates and 19 in LEA10 immunoprecipitates. (B) Name and description of the six common protein candidate interactors identified in LEA4, LEA10 and LEA5 immunoprecipitates are related to water deprivation. (C) GO Enrichment Analyses (GOEA) of the six common interactors identified in LEA4, LEA10 and LEA5 immunoprecipitates. (D) CNET plot showing the association of the six common interactors (green) to each GO category, denoted by edges. Red-blue scale corresponds to the adjusted p-value shown in C. (E) Clustered GOEA for Biological Process for all high confidence interactors found across immunoprecipitated LEAs: Among the interactors for all three LEAs there is an overrepresentation of water transport functions, whereas the partners specific for each LEA seems to support specialized functions.

### LEA5 interacts with water-deficit stress related proteins

Among the three immunoprecipitation experiments, the most successful in terms of the total number of candidate interactors was the immunoprecipitation of *pLEA5:LEA5-GFP* plants, with 81 candidate protein interactors of LEA5 identified (Supplementary Table 6). In agreement, LEA5 protein showed high accumulation levels in Arabidopsis roots in salt stress (Figure 2). Therefore, we decided to focus on the LEA5 candidate interactors to further investigate their functional relevance in water-deficit stress responses. First, the candidate interactors present in all the three LEA5 immunoprecipitation replicates were selected, resulting in 62 high confident candidate protein interactors (Supplementary Table 8). Next, eight candidate protein interactors of LEA5, including the bait protein LEA5 itself, were selected based on (i) their presence in all the three biological replicates (ii) their high fold change, indicating significant enrichment in the sample compared to the negative control, (iii) their presence in size exclusion chromatography (SEC) experiments in correlation with LEA5 (Veyel et al. 2018), (iv) the basal expression of their mRNA levels in root and stomata, and (v) their mRNA positive response to ABA and salt treatments, as reported in previous studies (Table 2). Among the eight selected candidate interactors, three were commonly detected in the three dehydrin protein complexes: LEA10, PlP2B and AHA1; while five of them were specific from the LEA5 protein complex: LEA4, LEA5, REDUCED HYPEROSMOLALITY-INDUCED [Ca^2+^] INCREASE 3 (OSCA3), PHOSPHOLIPASE D ALPHA 1 (PLD*α*1), and LEA8 (Table 2). Note that LEA4 fold change was below 1.5 (1.33) but still included for further analysis.

**Table 2.**
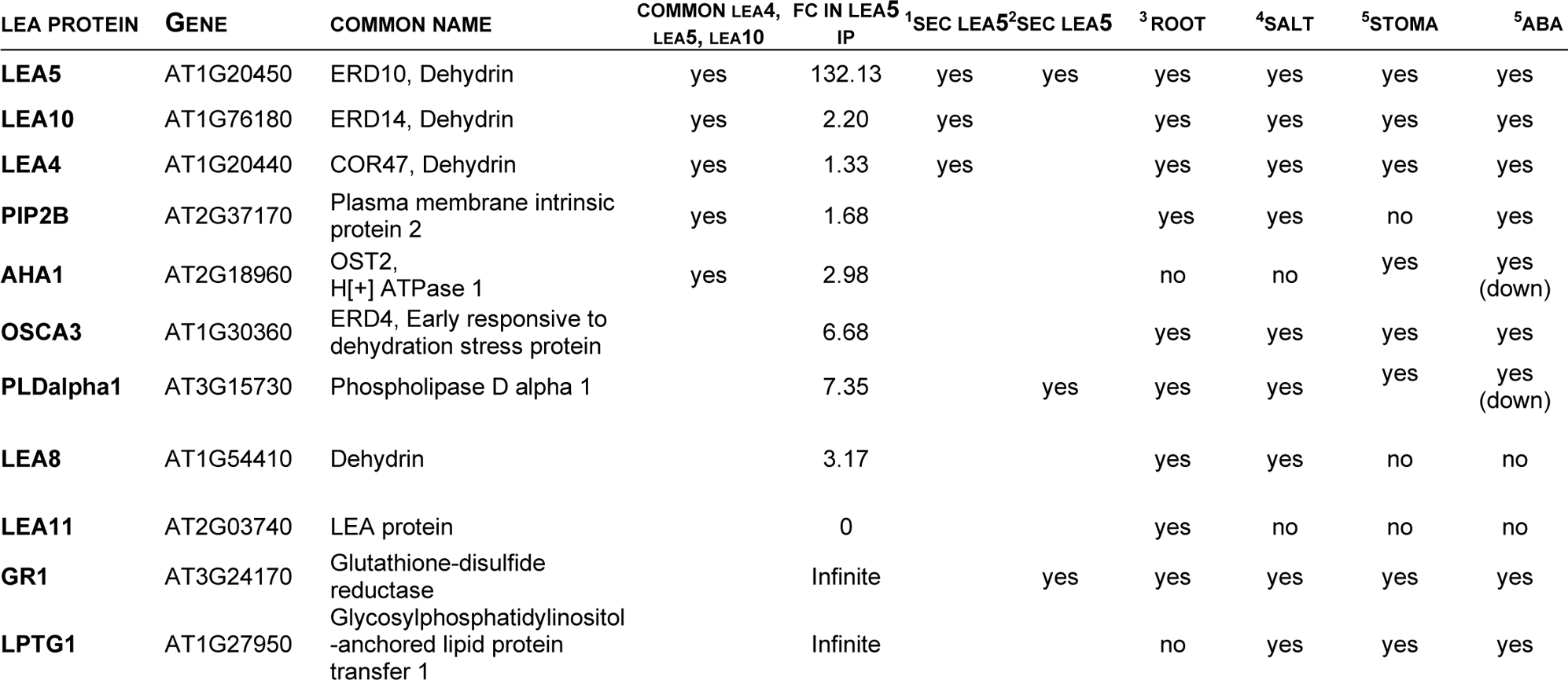
Selected LEA5 candidate protein interactors and their transcriptional response to stress. Data extracted from **^1^**SEC data with Pearson correlation > 0.7 from (Aryal et al. 2017); **^2^**SEC data from PROMIS (Veyel et al. 2018); **^3^**(Schmid et al. 2005); **^4^**(Kilian et al. 2007); **^5^**(Pandey et al. 2010

We aimed to validate direct protein-protein interactions of the LEA5 protein complex by using both Split Luciferase and ratiometric Bimolecular Fluorescence Complementation (rBiFC) assays. In addition to the eight candidate interactors selected above, we included three negative controls: two LEA5 candidate interactors that appeared only in two out of the three biological replicates: GLUTATHIONE-DISULFIDE REDUCTASE (GR1) and GLYCOSYLPHOSPHATIDYLINOSITOL-ANCHORED LIPID PROTEIN TRANSFER 1 (LPTG1), and one protein that was not detected in the LEA5 immunoprecipitates: the non-dehydrin LEA11 protein (Table 2). Therefore, a total of eleven cDNA sequences corresponding to the selected candidate interactors were cloned in different Gateway plasmids (for details, see Material and Methods). Split Luciferase assays in Arabidopsis seedlings confirmed the direct protein-protein interactions between LEA5 protein and six candidate interactors: LEA10, LEA4, OSCA3, AHA1, PIP2B and PLD*α*1 *in planta* (Figure 4A). No interaction was observed for LEA5 with the other four candidates: GR1, LPTG1, LEA8, and LEA11. Since GR1, LPTG1, and LEA11 were used as a negative control, these results validate the specificity of our assay. Moreover, a ratiometric bimolecular fluorescence complementation assay (rBiFC, Grefen and Blatt 2012) in *Nicotiana Benthamiana* plants confirmed the direct protein-protein interactions of LEA5 with LEA5 itself (homodimerization), and with the six previously confirmed interactors: LEA10, LEA4, OSCA3, AHA1, PIP2B and PLD*α*1 proteins, *in vivo* (Figure 4B, C). Confocal microscopy analysis of *N. Benthamiana* leaf cells transiently transfected with LEA5 bait protein carrying the C-terminal of YFP and the six candidate interactors carrying the N-terminal of YFP resulted in reconstitution of YFP fluorescence in the cytosol in all the cases analysed (Figure 4C). Quantification of the reconstitution YFP fluorescence and statistical analyses confirmed the direct protein interaction of LEA 5 with the six interactors: LEA10, LEA4, OSCA3, AHA1, PIP2B and PLD*α*1 *in planta*. LEA5 protein fused to the nYFP but not to the cYFP was used as a negative control showing no YFP auto-reconstitution (Figure 4B, C). Altogether, these results indicate that LEA5 dehydrin protein directly interacts with LEA5 itself, LEA10, LEA4, OSCA3, AHA1, PIP2B and PLD*α*1 proteins *in vivo*.

**Figure 4.**
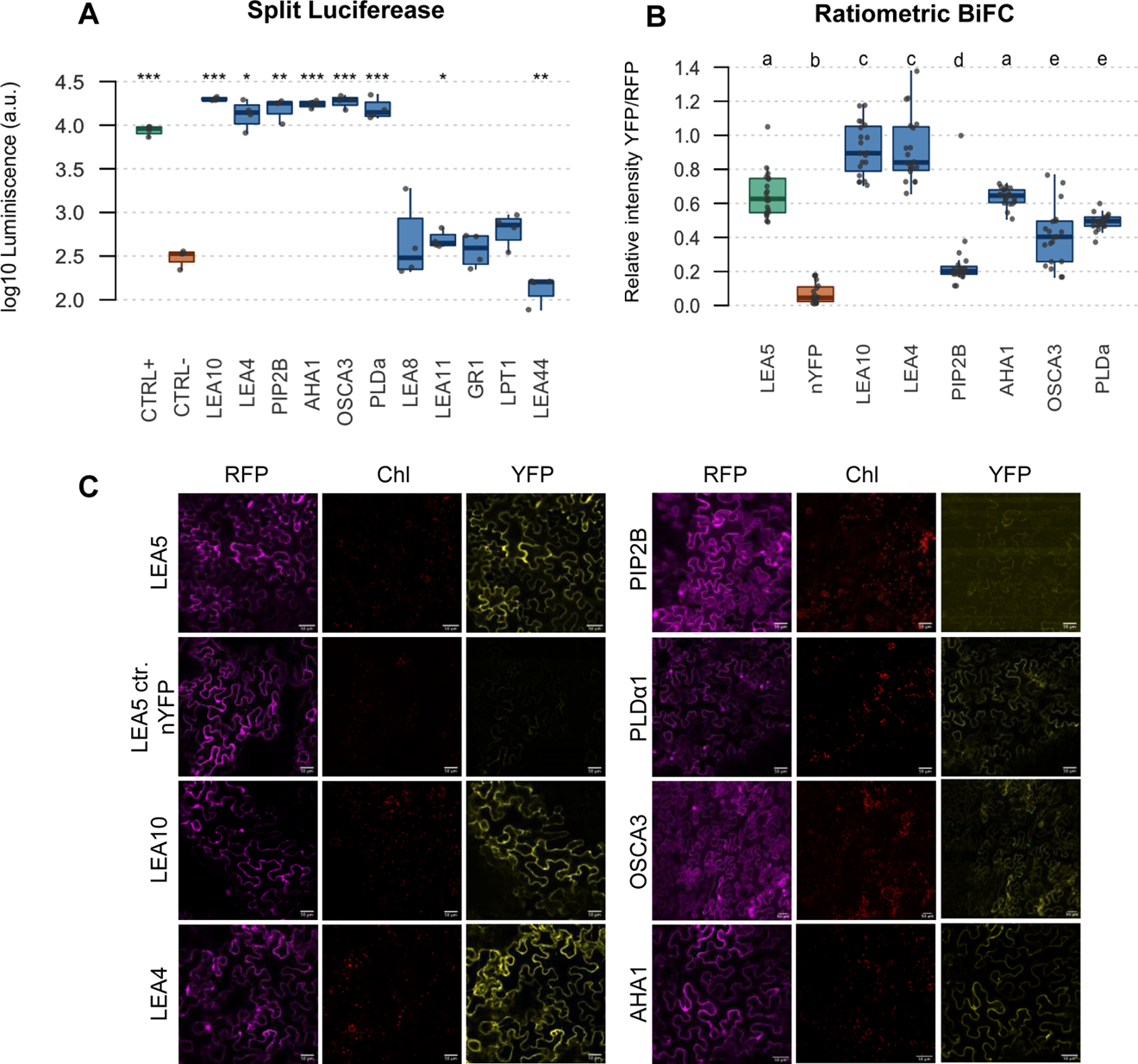
LEA5 high-confident candidates directly participate in protein-protein interactions. (A) Split-Luciferase complementation assays. Quantification of the reconstitution Luciferase luminescence and statistical analyses confirmed the direct protein interaction of LEA5 with the six candidate interactors: LEA10, LEA4, OSCA3, AHA1, PIP2B and PLD*α*1 *in planta.*. Asterisks represent p-values thresholds (*< 0.05, **< 0.01, ***<0.001) in a two-sided t-test pairwise comparison respect negative control (The bad quantile distribution in a one-way ANOVA restrained us to use this approach). (B) Ratiometric BiFC fluorescence quantification of the six candidate interactors: LEA10, LEA4, OSCA3, AHA1, PIP2B and PLD*α*1 *in planta*. The reconstituted YFP fluorescence is shown as proportional to the RFP cytoplasmic signal (YFP/RFP). LEA5 protein fused to the nYFP but not to the cYFP was used as a negative control showing no YFP auto-reconstitution. Relative fluorescence was analyzed using Image J. Data are means ± sem of 20 images selected at random over the surface of four-leaf samples in each case. Averages from three independent biological replicates (*n* > 31). Different letters represent significant differences (*p*-value < 0.05) in a one-way ANOVA plus Tukey’s HSD test. (C) Confocal microscopy analysis of *Nicotiana Benthamiana* leaf cells transiently transfected with LEA5 bait protein carrying the C-terminal and the five candidate interactors carrying the N-terminal half of yellow fluorescent protein (YFP). Confocal images acquired with a 20x objective. YFP, RFP and chlorophyll fluorescence channels. RFP was used as a control. LEA5 protein fused to the nYFP but not to the cYFP was used as a negative control showing no YFP auto-reconstitution. Scale bar: 50 µM

### LEA5 protein complex promotes seed germination and root growth under water-deficit stress conditions

To validate the functional relevance of the confirmed protein-protein interactions, two independent mutant lines for LEA5 and for each of the confirmed interactors were analysed (T-DNA insertion lines; Supplementary Table 9). Real time qPCR analyses revealed that at least one of the two lines showed significant lower mRNA expression levels than WT plants (Supplementary Figure 5). These were selected for further experiments. Double mutants were generated by crossing *lea5-1* loss-of-function mutant with the corresponding mutant of each confirmed interactor. However, no *lea5-1aha1-2* double mutant could be rescued in three independent attempts, and we argued that it might be lethal.

As salt stress induces *LEA5* gene expression and the protein complexes were characterized under salt stress, we first analysed the phenotypes of single mutants *lea5-1, osca3-2, pip2b-2* and *pldα1-2,* and double mutants *lea5-1osca3-2, lea5-1pip2b-2* and *lea5-1pldα1-2* germination in control or in salt stress conditions (Material and Methods). In one hand, neither single mutants of *LEA5*, *OSCA3*, *PIP2B* and *PLDα1* genes nor the double mutant combinations displayed any differences in terms of germination rates in four-day-old seedlings germinated in control conditions (Supplementary Figure 6A). However, *lea5-1* single mutants as well as *lea5-1osca3-2* and *lea5-1pip2b-2* double mutants showed a higher sensitivity to salt stress, as they all displayed a significant reduced germination rate when compared to WT seeds germinated in 150mM NaCl for four days (Figure 5A, B). In conclusion, these results show that LEA5 and the interaction of LEA5 with OSCA3 and PIP2B proteins promote seed germination under salt stress in Arabidopsis.

**Figure 5.**
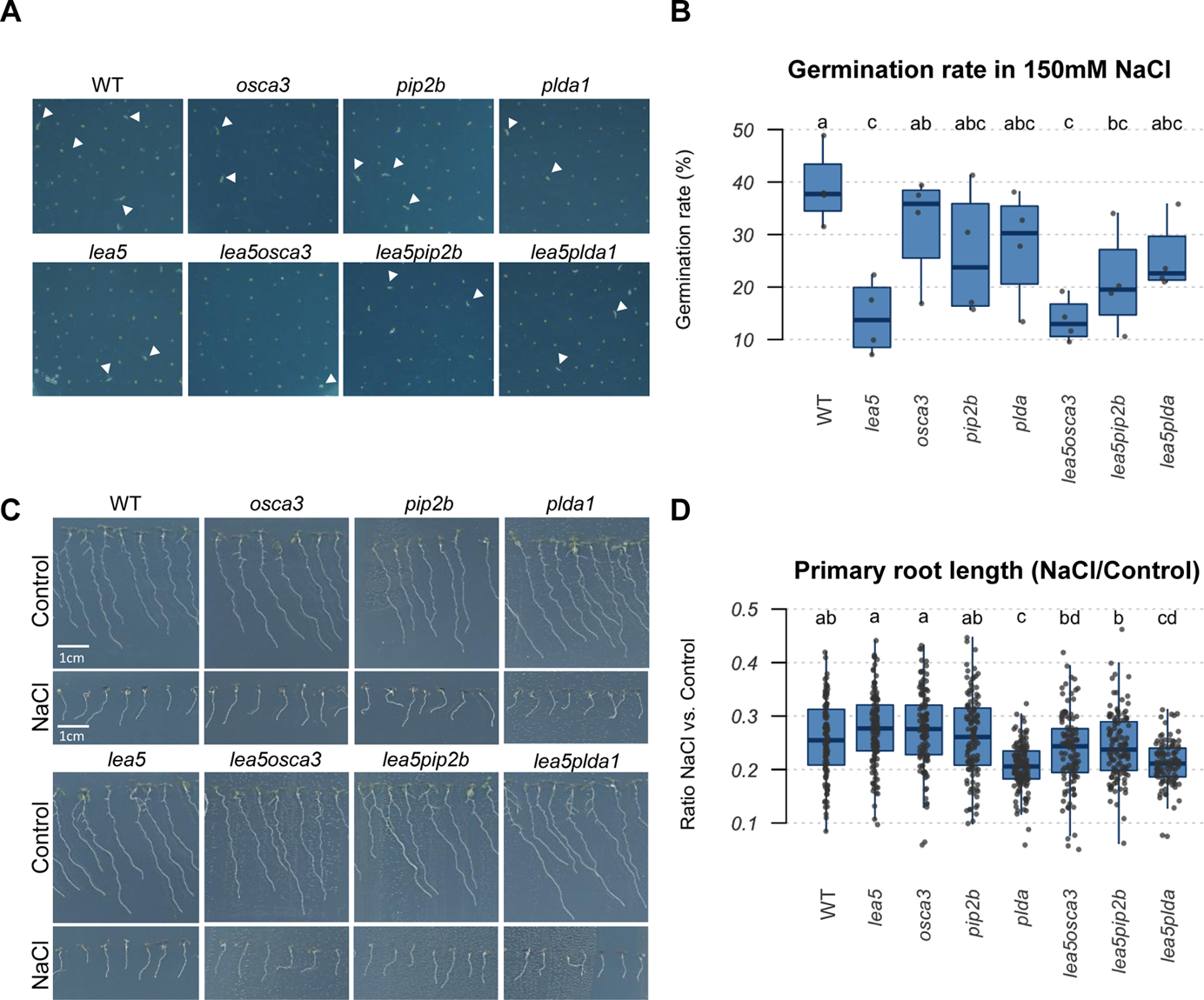
Salt stress negatively impacts in *lea5*, *lea5osca3* and *lea5pip2b* seed germination and in *pldα* and *lea5pldα* root length. (A) Representative images of Arabidopsis seeds after four days of germination in salt stress media with 150 mM NaCl. Examples of germinated seeds are indicated by white arrows. (B) Germination rates of Arabidopsis seeds after four days of germination in salt stress media with 150 mM NaCl. Data from four independent biological replicates were analysed. At least 50 seeds were quantified per genotype, and condition in each biological replicate. Shapiro-Wilk normality test revealed the normality of the data for each genotype and condition. Two-way ANOVA revealed a significant interaction between genotype and condition (p = 0.0013). Different letters represent significant differences in a one-way ANOVA plus Tukey’s HSD test (*p*-value < 0.05). (C) Representative images of eight-day old Arabidopsis seedlings in control conditions (upper panels) and in salt stress media with 150 mM NaCl (lower panels). (D) Root length sensitivity of eight-day old Arabidopsis seedlings after three days of growing in salt stress media with 150 mM NaCl. Boxplots depict the relative root length in NaCl conditions respect control. Ratios were calculated by dividing the measure root length seeds in salt stress by the median of the root length in control condition for each genotype and replicate. Data from four independent biological replicates were analysed. At least 20 roots were measured per genotype, and condition in each biological replicate. Shapiro-Wilk normality test revealed the normality of the data for each genotype and condition. Different letters represent significant differences in a one-way ANOVA plus Tukey’s HSD test (*p*-value < 0.05).

We also measured the root growth inhibition in salt stress conditions. For these experiments, plants were first germinated in normal conditions and transferred to salt stress media (Material and Methods). Neither single nor double mutants of LEA5 and its interactors displayed any differences in terms of root growth under control conditions (Supplementary Figure 6B). However, statistical analysis revealed that *pldα1-2* single mutants were sensitive to salt stress when compared to WT roots, and *lea5-1pldα1-2* double mutants showed an epistatic phenotype to *pldα1-2* parental lines (Figure 5C, D), indicating that LEA5 interaction with PLD*α*1 promote root growth under salt stress conditions. Since dehydrin proteins are also involved in other stress responses, we measured the root growth inhibition under osmotic stress conditions using mannitol (Material and Methods). *lea5-1pip2b-2* double mutants were more sensitive to osmotic stress-mediated root shortening when compared to WT and the corresponding parental single mutants *lea5-1* and *pip2b-2* in 300mM mannitol (Figure 6A, B). These results demonstrate that LEA5 interaction with PIP2B promotes root growth under osmotic stress.

**Figure 6.**
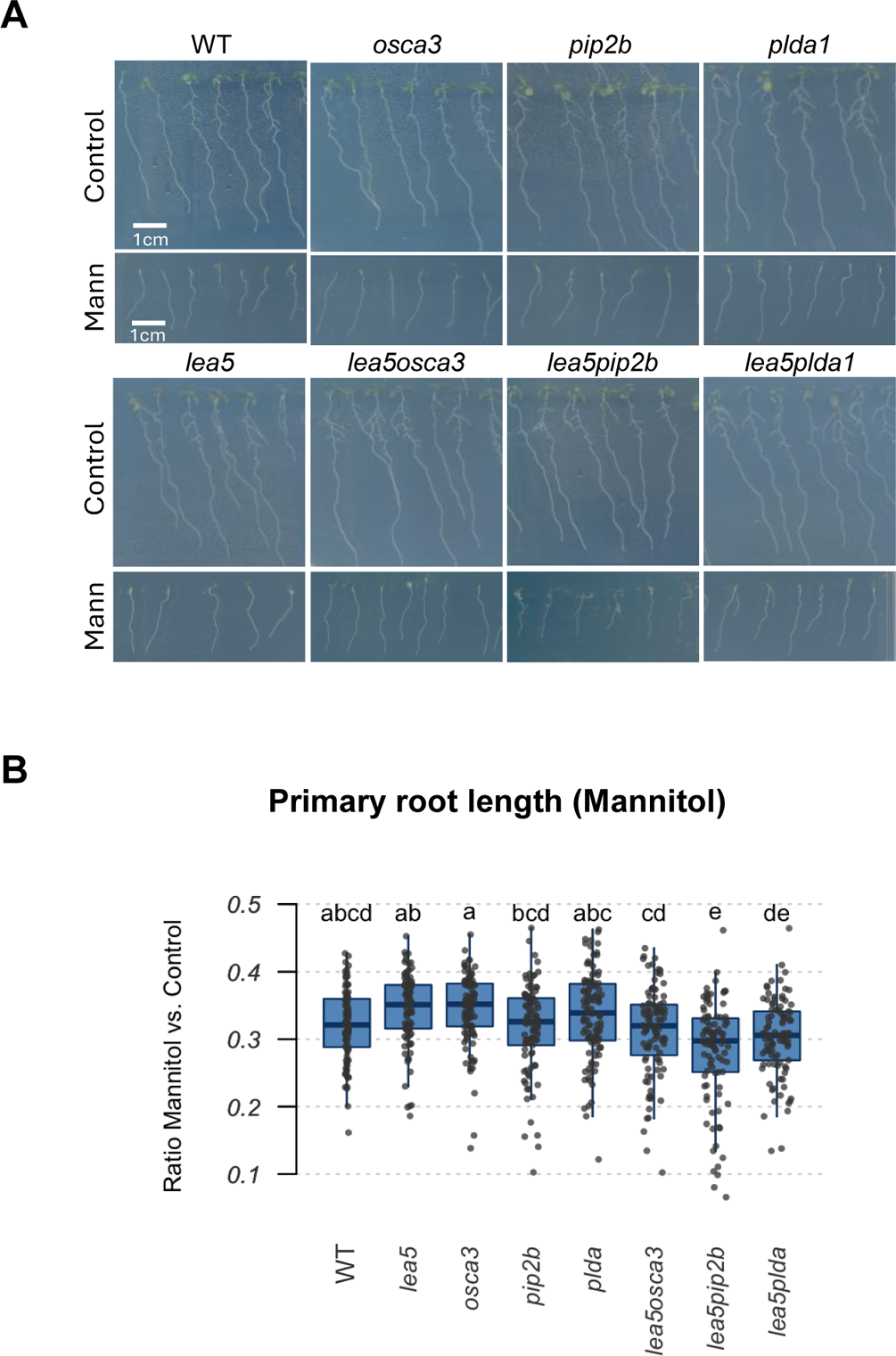
*lea5pip2b* roots are sensitive to osmotic stress. (A) Representative images of eight-day old Arabidopsis seedlings in control conditions (upper panels) and in osmotic stress media with 300 mM Mannitol (lower panels). (B) Root length sensitivity of Arabidopsis seedlings grown during three days in control media and transferred to osmotic stress media (300mM mannitol) for five days. Boxplots depict the relative root length in mannitol respect control. Ratios were calculated by dividing the measure root length seeds in osmotic stress by the median of the root length in control condition for each genotype and replicate. Data from five independent biological replicates were analysed. At least 20 roots were quantified per genotype, and condition in each biological replicate. Shapiro-Wilk normality test revealed the normality of the data for each genotype and condition. Different letters represent significant differences in a one-way ANOVA plus Tukey’s HSD test (*p*-value < 0.05).

In summary, our results show that LEA5 interacts with OSCA3, PIP2B and PLD*α*1 promoting plant germination and growth under salt stress conditions, while LEA5 and PIP2B interaction promotes root growth in osmotic stress conditions.

## DISCUSSION

While it is well reported that LEA proteins play a key role in water-deficit stress response and tolerance of plants and other organisms, the molecular mechanisms and pathways regulated by LEA proteins *in vivo* are not well understood. In this study, we report the molecular interactors of three LEA proteins from the dehydrin family in salt stress conditions. We have successfully identified novel *in vivo* protein complexes for the three dehydrin members LEA4, LEA5 and LEA10 under salt stress, we showed the direct interaction of LEA5 with LEA4, LEA10, AHA1, OSCA3, PIP2B and PLD*α*1 proteins, and we confirmed the functional relevance of LEA5 interactions with OSCA3, PIP2B and PLD*α*1 in promoting seed germination and root growth in high salinity and osmosis water-deficit stress conditions. All these results are summarized in a schematic model of the validated LEA5 protein interactions during salt stress reported in this study (Figure 7). A major strength of our study is the use *of in planta* immunoprecipitation from transgenic Arabidopsis lines expressing LEA proteins under their native promoters, a methodological advance that contrasts with previous studies relying on constitutive overexpression in plants and cell cultures or *in vitro* binding assays. This approach has enabled the capture of LEA interactors in their endogenous spatial and physiological context during stress.

**Figure 7.**
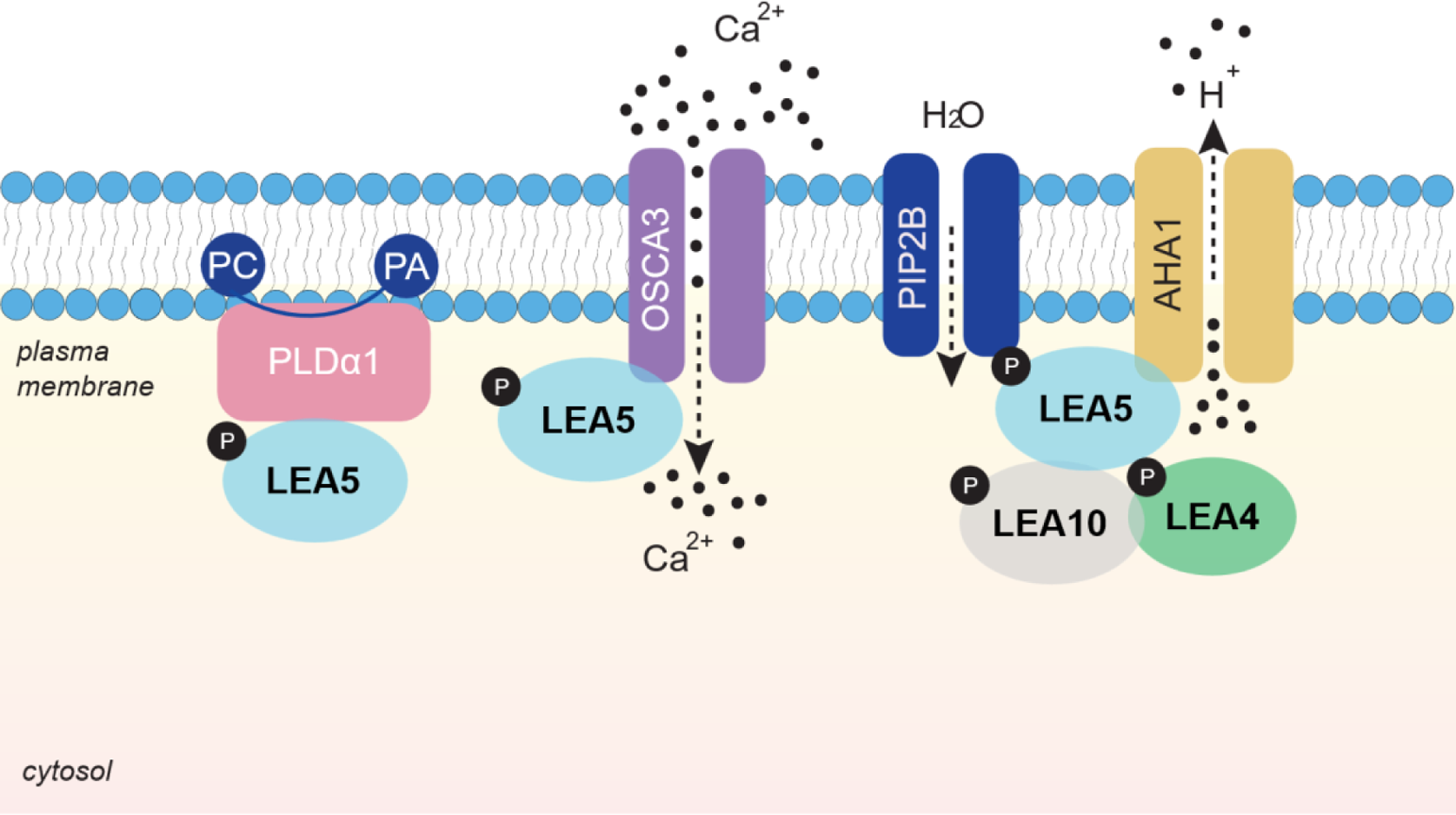
Model of validated LEA5 protein interactions during salt stress. The cartoon illustrates a schematic representation of the validated interactions of LEA5 protein during salt stress (24 h under 150 mM NaCl). The study demonstrates direct protein-protein interactions between the LEA5 dehydrin and LEA4, LEA10, PIP2B, AHA1, OSCA3 and PLDalpha1, validated through immunoprecipitation coupled to LC-MS/MS, split Luciferase, and BiFC assays. Interactions of LEA5 with PIP2B, OSCA3 and PLDalpha1 were further supported by functional genetic approaches. Abbreviations: PC, phosphatidylcholine; PA, phosphatidic acid; Ca^2+^, calcium ions; H^+^, hydrogen ions. The model assumes that LEA4, LEA5 and LEA10 dehydrin proteins undergo phosphorylation and folding upon stress, as previously demonstrated in vitro, enabling their interactions with other proteins.

### LEA *in vivo* immunoprecipitation approach

LEA proteins are highly disordered proteins, which makes them challenging for the study of their binding partners. To date, few molecular interactors have been reported for LEA proteins in Arabidopsis. Recently, tandem affinity purification (TAP) approaches using Arabidopsis cell cultures revealed LEA38 interaction with the chloroplast DEAD-box ATP-dependent RNA HELICASE 22 (RH22) (Karpinska et al. 2022) and phage display affinity experiments showed LEA14 dehydrin interaction with 35 potential client proteins (Unêda-Trevisoli et al. 2024). The latter study validated the direct protein-protein interactions of LEA14 with five client proteins through BiFC assays: PENTATRICOPEPTIDE REPEAT 596 (PPR596), SEED STORAGE ALBUMIN 4 (SESA4), RIBOSOMAL PROTEIN LARGE SUBUNIT 5A (RPL5A), and LEA14 itself, indicating homodimerization. However, the initial identification of LEA proteins candidate interactions in these studies relied only on *in vitro* approaches.

Here, we have successfully immunoprecipitated three LEA dehydrin protein complexes *in vivo*, as evidenced by the high enrichment of the bait protein abundance in each list of candidate interactors (Supplementary Tables 4, 5 and 6). Importantly, LEA proteins fused to GFP were expressed under their native promoters, and therefore the protein complexes were identified within the native cell types and tissues where LEA proteins are localized in the plant. Notably, our previous attempts to immunoprecipitate LEA proteins under control conditions consistently failed, suggesting that LEA proteins may remain largely unstructured and unfolded in the absence of stress, thereby limiting their capacity to form stable or specific protein–protein interactions. This observation aligns with the notion that LEA proteins gain functional conformation and binding capacity only upon cellular stress conditions that induce partial folding or structural ordering.

Moreover, the negative results of the direct interaction assay of LEA5 with GR1, LPTG1 and LEA8, underscore the importance of validating immunoprecipitation coupled to LC-MS/MS data with alternative approaches. In agreement with our results, previous studies identified LEA8 in monomeric forms and not interacting with other proteins in Arabidopsis (Aryal et al. 2017). In our study, GR1 and LPTG1 were identified only in two out of three biological replicates of the LEA5 immunoprecipitation experiments, indicating that the presence of a candidate interactor in all the biological replicates might be an important filter to eliminate false-positive interactors in protein–protein interaction studies. Alternatively, these results could also indicate that these proteins are in the same protein complex than LEA5 yet do not interact directly with the bait dehydrin LEA5 protein.

### LEA5 interaction with LEA and other proteins

Since LEA5 immunoprecipitation experiments were the most successful in this study, eight high-confidence LEA5 candidate interactors were selected to further investigation of their functional relevance in water-deficit stress responses. However, only five LEA5 interactors — LEA4, LEA5, LEA10, PlP2B, OSCA3, and PLDα1— were validated both at the biochemical and genetic levels.

First, we observed that LEA4, LEA5 and LEA10 dehydrin proteins heterodimerize with each other in response to salt stress. We identified LEA5 and LEA10 in LEA4 immunoprecipitates, LEA4 and LEA10 in LEA5 immunoprecipitates, and LEA4 in LEA10 immunoprecipitates (Supplementary Tables 4, 5 and 6). Our semiquantitative rBiFC assays confirmed a strong and direct interaction of LEA5 with LEA5, LEA4 and LEA10 (Figure 4). Moreover, the LEA4_1 outlier replicate sample that was removed, showed LEA5 and LEA10 as the only two candidate proteins with a FC above 1.5. This supports the notion that heterodimerization among the three LEA proteins is highly stable, even under compromised experimental conditions, as observed in this replicate. In agreement, previous global analysis of plant protein complexes using SEC coupled with high-resolution LC-MS/MS analysis reporting cytosolic protein complexes detected the oligomerization of these three dehydrin proteins (Aryal et al. 2017). Also supporting these results, previous BiFC experiments showed that three dehydrin proteins LEA4, LEA5 and LEA51 homodimerize and heterodimerize in different cell compartments (Hernández-Sánchez et al. 2017). At the genomics level, the three dehydrin genes *LEA4*, *LEA5* and *LEA10* cluster together (Hundertmark and Hincha 2008) and while they all lack the Y-segment, they carry the more recently described F-segment (Strimbeck 2017). A large-scale genomic analysis of 60 different organisms identified two different communities of dehydrin genes (Artur et al. 2019). The study suggests that phylogenetic separation of these communities occurred in angiosperms and report that dehydrin genes encoding proteins from community 1 (LEA33, LEA34, LEA44, LEA45) are primarily expressed during seed development, while genes encoding proteins from community 2 (LEA4, LEA5, LEA10, LEA44) appear to be specifically induced by abiotic stresses like drought, heat, and salinity (Artur et al. 2019). Three dehydrin proteins from the community 1, LEA4, LEA5, and LEA10 are phosphorylated *in vivo* by the Casein Kinase 2 in Arabidopsis (Alsheikh et al. 2003), and this phosphorylation regulates binding of the three dehydrin proteins to calcium ions *in vivo* (Alsheikh et al. 2003, 2005). Further studies should address whether phosphorylation of these dehydrins upon stress is critical for the organization of this core protein complex. In addition, nine common candidate interactors were identified in the three dehydrin LEA protein complexes. GO enrichment analyses revealed they are all involved in water-deficit stress related functions. Thus, it is tempting to propose that LEA4, LEA5 and LEA10 interact with these target proteins to stabilize them during stress. Altogether, these data suggest that LEA4, LEA5 and LEA10 form a core protein complex that likely play a primarily redundant or synergistic function in response to water-deficit stress responses. However, specific interactors were detected in LEA4, LEA5 and LEA10 protein complexes as well, which were enriched in distinct GO categories, suggesting that LEA interactor partners may take on more specialized functions while preserving a water stress response core.

Second, we report the direct interaction between LEA5 and the calcium channel OSCA3 at both the biochemical and the genetic levels, suggesting that LEA5 plays a role in calcium-mediated responses to water-deficit stress. Previous studies showed that LEA5 dehydrin is phosphorylated in response to osmotic stress and can bind to calcium ions (Alsheikh et al. 2005) suggesting that LEA5 might function as a calcium-dependent chaperone. OSCA1.1, OSCA1.2 and OSCA3.1 are structurally related members of the OSCA/TMEM63 family of mechanosensitive ion channels in Arabidopsis, all forming homodimers with a similar overall architecture (Murthy et al. 2018; Zhang et al. 2018). OSCA1.1 has been reported to function as an osmosensor mediating osmotic stress-induced Ca^2+^ increases in Arabidopsis (Yuan et al. 2014) and its crystal structure has been previously reported (Liu et al. 2018). More recently, the crystal structure of OSCA3.1 revealed a similar architecture to OSCA1.1 and OSCA1.2 (Jojoa-Cruz et al. 2024). While only OSCA1.1 has been reported to act as osmosensor (Yuan et al. 2014), it is tempting to propose that OSCA3 likely participates in water-deficit stress signaling. Supporting this hypothesis, overexpression of the closest homolog of OSCA3 in the extreme halophyte *Salicornia brachiate* in Arabidopsis led to an enhanced response to severe salt treatment (Jha and Mishra 2022). These transgenic plants exhibited enhanced germination and root length under salt stress, a better response to withholding watering and salt watering in soil, and increased accumulation of osmoprotectant metabolites such as sugars and proline (Jha and Mishra 2022). Similarly, the *Zea mays* orthologue of OSCA3, ZmERD4, is induced by drought, cold and salt stress, and the overexpression of ZmERD4 in Arabidopsis improved plants resistance to drought and salt stress (Liu et al. 2009). In our study, the *lea5-1* loss-of-function mutant showed hypersensitivity to salt stress in seed germination assays. Consistently, *lea5* mutants also exhibit increased sensitivity to cold stress and drought stress assays in soil (Kim and Nam 2010). While *osca3-2* single mutants did not show a phenotype in salt stress germination assays, the *lea5-1osca3-2* double mutant showed a hypersensitive phenotype that was epistatic to the *lea5-1* single mutant parental line. Because the *lea5-1osca3-2* double mutant phenotype is indistinguishable from the *lea5-1* single mutant, this indicates that the loss of LEA5 masks the effect of OSCA3, implying that LEA5 functions downstream of OSCA3 within the same signaling pathway rather than in a parallel, independent route. Altogether, these findings support a model in which LEA5 interaction with OSCA3 is involved in promoting seed germination under salt stress conditions by mediating calcium-dependent responses downstream of OSCA3 signaling pathways.

Third, LEA5 interacts with PIP2B aquaporin. Specifically, we identified PIP2B as a common interactor or LEA4, LEA5 and LEA10 dehydrins. It was previously shown that PIP2B interacts with LEA4, LEA5 and LEA51 proteins *in vivo* through BiFC assays (Hernández-Sánchez et al. 2019). The authors proposed that interactions among dehydrins and PIP2B may directly prevent aquaporin denaturation and inactivation. PIP2B has been shown to be degraded in drought stress conditions (Lee et al. 2009). Here, we confirmed the direct interaction of LEA5 with PIP2B in salt stress conditions, suggesting that LEA5 is likely protecting PIP2B from stress-mediated degradation. While *pip2b-2* single mutants did not show a phenotype in salt stress germination when compared to WT plants, *lea5-1pip2b-2* double mutants showed more sensitivity than WT plants in salt stress germination, which is epistatic to the *lea5-1* parental line. However, the phenotype of the double mutant *lea5-1pip2b-2* is weaker than the phenotype of the double mutant *lea5-1osca3-2* plants. In agreement, rBiFC results show a weaker interaction of LEA5-PIP2B compared to other confirmed interactors (Figure 4). Moreover, as PIP2B has been identified as a common interactor in the three dehydrin complexes, it is tempting to propose that LEA4, LEA4 and LEA10 are playing a redundant role protecting PIP2B protein from stress-mediated degradation. Finally, *lea5-1pip2b-2* double mutants showed more sensitivity than WT roots under mannitol treatment, suggesting that this interaction is also functional in other osmotic stress conditions. Altogether, these results suggest that LEA5 and PIP2B interact at the biochemical and functional levels to promote seed germination under water-deficit stress conditions. A recent work identified SYNTAXIN OF PLANTS 132 (SYP132) together with the plasma membrane ATPase AHA1 and PIP2B in the immunoprecipitates of PIP2A protein under salinity stress. Within the PIP2A candidates’ interactors OSCA3 was also identified, among others (Baena et al. 2024).These results are in agreement with the elucidation of LEA5 protein complex under salt stress, containing AHA1, PIP2B and OSCA3, among others, that we have identified in this study.

Lastly, we identified a direct biochemical interaction between PLDα1 and LEA5 under salt stress conditions. Consistent with previous findings showing sensitivity of the *pldα1* mutant allele SALK_053785 during germination under salt stress (Hunter et al. 2019), the *pldα1-2* single mutants (SALKseq_072461.2) also displayed reduced root growth in response to salt stress, yet we did not observe a significant phenotype for neither *pldα1-2* nor *lea5-1pldα1-2* in seed germination assays. Notably, in our assays, *lea5-1pldα1-2* double mutants showed a salt-sensitive root growth phenotype that was epistatic to the *pldα1-2* parental line while *lea5-1* parental line was indistinguishable from WT plants. This genetic relationship suggests that LEA5 and PLDα1 act in the same pathway. Because the double mutant did not revert the phenotype observed in the *pldα1-2* single mutant, it is likely that PLDα1 functions downstream of LEA5 during salt stress response. These findings support the hypothesis that LEA5 and PLDα1 interact both biochemically and functionally to promote root growth under salt stress conditions. In line with this, CYS-RICH RECEPTOR-LIKE KINASE 2 (CRK2) has been reported to enhance salt stress tolerance and to physically interact with PLDα1 and PIP2B, among other proteins (Hunter et al. 2019). Corroborating this interaction network, we also identified PIP2B as a LEA5 interactor, reinforcing their plausible role as a protein complex in water-deficit stress responses.

Moreover, PLDα1 is phosphorylated in response ABA treatments (Wang et al. 2013) and plays a pivotal role in ABA-mediated stomatal closure (Uraji et al. 2012). *pldα1* loss-of-function mutants (SALK_053785) and PLDα1-antisense lines exhibit increased water loss compared to WT plants. Interestingly, stomatal closure in *pldα1* mutants is unresponsive to ABA but can still be induced by phosphatidic acid (PA), suggesting a bypass mechanism (Zhang et al. 2004). In wild-type plants, ABA triggers PA production from phosphatidylcholine via PLDα1, a response absent in *pldα1* mutants. This PLDα1-derived PA binds to the PP2C phosphatase ABI1 and modulates ABA signaling (Zhang et al. 2004). LEA5 protein accumulates in response to ABA in our study (Figure 2), indicating that its interaction with PLDα1 may form part of an ABA-dependent response induced by salt stress. Previous immunoprecipitation experiments with the ABA-dependent subclass III SnRK2.2 (Soma et al. 2023) and SnRK2.6 (Waadt et al. 2015), revealed interactions between these two kinases. In our analysis, we have identified 26 candidate interactors shared between LEA5 (this study) and SnRK2.2 (Soma et al. 2023), while only two candidate interactor overlaps between LEA5 and SnRK2.6, namely the copper chaperone AT3G56240 and the LEA8 protein AT1G54410. However, PA is also known to bind to subclass I SnRK2 kinases, such as SnRK2.4, which function independently of ABA (Julkowska et al. 2015). In addition, LEA5, LEA10, REMORIN1.2, REMORIN1.3 and HSP70-interacting protein 1 were also identified as phosphorylation targets of the subclass I SnRK2.10 (Maszkowska et al. 2019). All of these were also found among the LEA5 candidate interactors in our immunoprecipitation experiments. Moreover, LEA5 protein is more accumulated in response to salt stress than to ABA as shown in Figure 2. Altogether, these results suggest that LEA5 is also phosphorylated and activated through an ABA-independent mechanism. Taken together, these findings suggest that LEA5 is involved in both ABA-dependent and ABA-independent signaling pathways in response to salt stress.

Our initial hypothesis proposed that LEA proteins function as hub molecules linking signalling and metabolic pathways. In this study, several candidate interactors of dehydrin proteins, including metabolic enzymes such as glutathione and dehydroascorbate reductases, ATPases, calcium and water channels, and phosphatases such as RCN1 PP2A, among others, were successfully detected. However, we did not identify any kinase signaling proteins as a candidate upstream interactor of the dehydrins proteins, which could potentially phosphorylate them in response to water-deficit stress. We argue that this is because we performed the immunoprecipitation experiment after 24 hours of salt stress treatment while protein phosphorylation upon water-deficit stress occurs within minutes (Lin et al. 2020; Soma et al. 2020). Supporting this interpretation, a phosphoproteomics study identified LEA5 and LEA10 as phosphorylation targets of the ABA-independent subclass I SnRK2 kinase, SnRK2.10, after 30 minutes of salt stress treatment (Maszkowska et al. 2019). Future studies focusing on early timepoints will be essential to capture transient signaling components upstream of LEA proteins under stress.

Furthermore, in the current study, we observed an enrichment in plasma membrane proteins in the LEA5 and LEA10 immunoprecipitates, despite both proteins are localized in the cytosol according to previous studies (Candat et al. 2014). In agreement with our findings, previous studies identified LEA4, LEA5 and LEA10 as putative plasma membrane proteins (Kawamura and Uemura 2003), demonstrated that dehydrins bind to lipid vesicles *in vitro* (Ismail et al. 1999; Hara et al. 2001; Koag et al. 2003, 2009) and showed that the K-segment is necessary for liposome binding (Eriksson et al. 2011, 2016). The interaction of LEA proteins with model membranes has been explored *in vitro* (Thalhammer et al. 2014; Atkinson et al. 2016; Eriksson et al. 2016; Singh and Graether 2021). Moreover, LEA protein interactions with the plasma membrane are often associated with folding upon binding (Rahman et al. 2013; Atkinson et al. 2016; Eriksson et al. 2016; Singh and Graether 2021). In agreement with these results, F-segment dehydrins have been described to play potential functional roles in membrane and protein binding (Richard Strimbeck 2017). Therefore, these results indicate that dehydrin LEA proteins functions are highly related to the plasma membrane. Taken together, our results point to a prominent role for LEA proteins at the membrane interface, where they may stabilize protein complexes and lipids under stress.

Finally, while we report a promoting role for LEA5 dehydrin in seed germination under salt stress, a recent study shows an inhibitory role for dehydrin LEA14 in seedling germination under salt stress (Perrella et al. 2024). The study shows that LEA14 and MADS AFFECTING FLOWERING 5 (MAF5) inhibit seedling establishment under salt stress downstream of the HISTONE DEACETYLASE COMPLEX 1 (HDC1). HDC1 attenuates transcriptome re-programming in salt-treated seedlings. In conclusion, even though dehydrins are activated in response to stress, the function of each member of the family in the plant is still poorly understood and is likely specialized to different types of stress and tissues. Moreover, the identification of nine conserved protein interactors across LEA4, LEA5, and LEA10 in our study further supports that LEA proteins engage in specific molecular interactions with their target interactors rather than binding promiscuously. Collectively, our findings underscore that LEA proteins execute specialized, non-redundant roles in the plant response to water-deficit stress, challenging the long-standing assumption that their function is primarily structural or non-specific.

## CONCLUDING REMARKS

The identification of interacting proteins of dehydrin and other LEA proteins is crucial for unravelling the molecular responses of plants in water-deficit stress. In this study, we successfully performed *in vivo* protein complex immunoprecipitations of the LEA4, LEA5 and LEA10 dehydrin proteins, identified the candidate interactors using LC-MS/MS, and confirmed specific protein-protein interactions with LEA5 using Split-luciferase and BiFC analyses. Our major findings are: (i) LEA5 forms homo- and hetero-oligomers with the other dehydrins LEA4 and LEA10 (Figure 7); (ii) LEA4, LEA5 and LEA10 dehydrins share a common core of proteins related to water deficit stress; (iii) LEA5 interacts with well-described water-deficit stress related proteins such as aquaporins (PIP2B), calcium channels (OSCA3), and lipid metabolism enzymes (PLD*α*1) (Figure 7); (iv) Dehydrin candidate interactors are enriched in plasma membrane proteins; (v) LEA5 interacts with OSCA3, PIP2B and PLD*α*1 to promote root growth and seed germination in water-deficit stress conditions.

In conclusion, we provide novel evidence of the *in vivo* targets of dehydrin LEA proteins under salt stress, and functional insights into the roles of LEA5 dehydrin protein complex in salt and osmotic water-deficit stress responses. Altogether, these results challenge the traditional view of LEA proteins as non-specific structural protectants and uncover their roles as dynamic interactors with key stress-related proteins. Further understanding the specific roles of the different LEA proteins *in vivo* will contribute to elucidating signaling and molecular mechanisms involved in plant stress adaptation.

## MATERIAL AND METHODS

### Plant growth

Arabidopsis seeds were sterilized with 0.2% sodium hypochlorite for 5 min, rinsed with water four times and vernalized for 48 h in dark conditions and at 4 °C. Seeds were sowed in *in vitro* plates filled with half-strength Murashige and Skoog media (MS) and grown for 7 days in long-day conditions (16 h light/8 h dark) at 22 °C. Then, 7-day-old seedlings were transferred to pots containing approximately 30 g of universal substrate supplemented with perlite and vermiculite. Plants were let growth in a chamber with long day conditions at 22 °C and a relative humidity of 60%.

### Translational reporter lines

To generate the LEA native promoter translational transgenic lines (*pLEA:LEA-GFP*) up to 2kb of the *LEA* promoter sequences and the full-length genome sequence of *LEA* genes (including introns, excluding the last stop codon) were amplified by PCR using specific primers (Supplementary Table 9) from genomic DNA of Arabidopsis Col-0 plants. The resulting PCR products were cloned into the entry vector pDONR201 by recombination using BP Clonase II Enzyme mix (Invitrogen Technologies). The resulting entry clones were sequenced and subsequently recombined into a pGWB604 destination vector (R1-R2-sGFP-Tnos) using Gateway LR Clonase II Enzyme mix (Invitrogen Technologies). Cultures of Agrobacterium tumefaciens GV3101 were transformed, and transgenic plants were generated by floral dipping into Arabidopsis Col-0 plants background as previously described (Clough and Bent 1998). In the third generation, two independent homozygous lines were selected for each LEA protein.

### Isolation and confirmation of *knock-out* mutants

Seeds for each T-DNA insertion line were ordered from the corresponding NASC and GABI seedstocks centres. All SALK and GABI insertional lines were genotyped using their corresponding specific primers described in Supplementary Table 9. For gene expression analysis, total RNA was extracted from Arabidopsis seedlings using Mini spin kit for RNA isolation from plant (Macherey-Nagel). RNA was converted to cDNA by reverse transcription using First Strand cDNA Synthesis kit (Roche). RT qPCR was done using specific F and R primers for each studied gene for *UBQ30* as a housekeeping gene. Transcript levels were analysed by RT qPCR using SYBR Green master mix (Applied Biosystems™).

### Water-deficit stress responses phenotypic assays

For phenotypic analysis, Arabidopsis seeds were sterilized with 0.2% sodium hypochlorite for 5 min, rinsed with water four times and stratified 24 hours in dark conditions at 4 °C. For determining seed germination in salt stress assays, at least 100 per genotype surface sterilized seeds from Col-0 WT, single and double loss-of-function mutants were sowed in half strength Murashige and Skoog (MS) medium containing 0, 100, 150, 200 and 250 mM NaCl, 1% (w/v) plant agar, 1% (w/v) sucrose each and grew in a controlled chamber in long-day conditions (16 h light/8 h dark) at 22 °C. Pictures were taken every day from 3 to 7 days after sowing to determine the germination rate. For determining root length in salt and osmotic stress assays analysis, Arabidopsis seeds were sterilized as described above. At least 40 seeds per genotype were sowed in half-strength MS media containing 1% (w/v) plant agar and 1% (w/v) sucrose and grown. The plates were maintained vertically for 3 days in long-day conditions (16 h light/8 h dark) at 22 °C. After that around 20 plants were transferred to half strength half-strength MS media (control), and other 20 plants were transferred to media supplemented with 150mM of NaCl or 300mM of mannitol (treatment) and grown for an additional 5 days. Pictures were taken with a flatbed scanner, and roots were measured using ImageJ.

### Protein extraction and western blot

Total protein content was extracted by homogenizing nitrogen-grinded seedlings in two volumes of extraction buffer (50 mM Tris-HCl pH 7.5; 150 mM NaCl; 1% Triton X-100 and 1x complete protease inhibitor pill (Roche) for each 100ml of buffer) on ice. Extracts were incubated 10 min on ice, homogenized by 5 sec vortex and centrifuged 30 min at 13000 rpm at 4 °C. Supernatants were used to assess protein concentration using by Pierce™ BCA Protein Assay Kits (Thermo Fisher Scientific). A total of 60 ug of protein was loaded in each well. Protein samples were prepared by adding loading buffer (80mM Tris-HCl pH 6.8; SDS 1.6%; DTT 0,1M; Glycerol 5%; Bromophenol Blue) and denatured by boiling samples at 95°C for 5 minutes. Samples were run in an SDS-PAGE gel for 1 h 30 min and transferred by blotting at 100 mV for 1 h into nitrocellulose membranes (Hybond-ECL, GE Healthcare®). Membranes were incubated with antiGFP-HRSP antibodies (MACS, Miltenyi Biotech) for 1 h and revealed with Pierce™ Western Blot Detection Kit (Thermo Fisher Scientific) in a ChemiDoc chemiluminescence imaging system (Bio-rad) for 1 minute exposure.

### Protein extraction and *in vivo* immunoprecipitation

10-day-old Arabidopsis seedlings were harvested and crosslinked by vacuum infiltration in a 1% formaldehyde in ice-cold PBS buffer for 30 min (Hall and Struhl 2002). The reaction was quenched by adding cold 200 nM glycine for 30 min. Cross linked tissue was then washed with PBS and stored at −80 °C. Samples were grinded manually in a pre-chilled mortar in liquid nitrogen. The protein extraction (50 mM Tris-HCl pH 7.5; 150 mM NaCl; 1% Triton X-100; 1x complete protease inhibitor pill (Roche) for 100 ml of buffer) and washing buffers (50 mM Tris-HCl pH 7.5; 150 mM NaCl; 0.1% Triton X-100; 1x complete protease inhibitor pill (Roche) for 100 ml of buffer) were freshly prepared and chilled on ice before starting extraction. Powdered tissue was resuspended with the extraction buffer using a 1:2.5 ratio (10 ml fresh tissue in 25 ml of extraction buffer) and incubated 10 min on ice. Protein extracts were transferred into a glass plunger tube and homogenized by moving the plunger up and down 10 times. The total homogenised protein extracts were transferred into new 50 ml tubes and incubated on ice for 20 min. The lysates were centrifuged in a precooled centrifuge at 20000 *g* for 10 min at 4°C. The supernatants (soluble protein extracts) were transferred to new centrifuge tubes and spin at 20000 g for 10 min at 4 °C (x2). The final protein extracts were transferred into new 50ml tubes and 200 μl of anti-GFP microbeads were added to each sample and incubated for 1 h on a rotating device at 4 °C. μ-Columns were placed in the μMACS separator and calibrated with 200 μl wash buffer. The immunoprecipitates were applied onto the columns and let them run through by gravity flow. 50μl of flow through were aliquoted in SDS loading buffer and stored at −80 °C as not bound flow through sample for the western blot. The immobilized beads were washed with 200 μl of wash buffer (4 x). Freshly prepared 20 μl of 8 M urea were applied onto the columns and incubated for 10 min at room temperature (20 °C). Immunoprecipitated samples were eluted with 50 μl of 8 M urea into a fresh 1.5 ml Eppendorf low-protein-binding tube. The protein immunoprecipitated sample was stored at −80 °C.

### FASP digestion and sample preparation for LC-MS/MS analysis

The protein immunoprecipitated samples were defrost on ice and 50 μl of sample were mixed with 100 µl 8 M urea in 10 mM Tris-HCl, pH 8 and applied to Filter-aided sample preparation (FASP) filter columns, and centrifuged at 10000 *g*, 4 °C, 5min. Note: It is recommended maximum of 250 µg total protein. A rough approximate quantification of protein concentration by dropping 1 µl of BSA curve (above: from 2 µg/µl to 30 ng/µl) and 1 µl of each protein extraction sample was used. 100 µl of 8 M urea in 10 mM Tris-HCl, pH 8 were added in the immunoprecipitate tube to clean remaining protein sample and applied to the filter. Filters were centrifuged at 14000xg, 40min at RT, and flow-through was discarded. 50 µl of 10 mM DTT in 8 M urea/10 mM Tris-HCl, pH 8 were added. Filters were centrifuged at 14000 *g*, 30 min at RT, and flow-through was discarded. 50 µl of 27 mM iodoacetamide in 8 M urea/10 mM Tris-HCl, pH 8 were added and filters were sealed with aluminium and mixed at 600 rpm in thermo-mixer for 1 min and incubated without mixing for 5 min. Filters were centrifuged at 14000 *g*, 30 min at RT, and flow-through was discarded. 100 µl of 8 M urea/10 mM Tris-HCl, pH 8 were added and filters were centrifuged at 14000 *g* for 40 min at RT, and flow-through was discarded. The filter units were transferred into a new collection tube. 20 µg of Sequencing

Grade Modified Trypsin were diluted in 1 ml of buffer (Promega; V5111; 5 x 20 µg) obtaining 0,02 µg/µl Trypsin. One Unit of trypsin enzyme was added for each 25 Unit of protein. Tubes were covered with parafilm, wet paper and aluminium foil, placed in a wet chamber and incubated at 37°C overnight. Day after, samples were centrifuged at 14.000 *g* for 40 min at RT. 50 µl of NaCl 0.5 M were added and centrifuged at 14.000 g for 20 min at RT to elute the peptides. 15 µl of Trifluoroacetic acid (TFA) 10% were added to have TFA at 1% final concentration in the eluted peptides samples. pH was checked with a pH indicator strip to ensure pH was below 3. Seppack Columns were used under vacuum conditions and activated with 1 ml of 100% methanol followed by 1 ml of 80% Acetonitrile (ACN) in 0.1% TFA. Columns were equilibrated with 0.1% TFA (x2). 200 µl 0.1%TFA were added in the peptides samples and samples were loaded into the Seppack Columns. 200 µl of 0.1% TFA were added to clean the remaining samples in the eppendorf and loaded into the column. Samples retained in the column were washed with 1 ml 0.1 % TFA (x2). Peptides were eluted with 800 µl of 60% ACN in 0.1% TFA and dried in the Speed vacuum for at least 5 hours. Peptides were stored at −80°C.

### LC-MS/MS and protein identification and quantification

Dried peptides were resuspended in MS loading buffer (3% ACN, 0.1 % FA) and measured with Q Exactive HF (Thermo Fisher Scientific) coupled to a reverse-phase nano liquid chromatography ACQUITY UPLC M-Class system (Waters). The gradient ramped from 5% ACN to 24% ACN over 6 min, then to 36% ACN over the next 10 min followed by a 2 min washout with 85% ACN. Data were acquired using a data-dependent MS/MS method (DDA) with the following settings: full scans were acquired at a resolution of 60 000, AGC target of 3e6, maximum injection time of 50 ms, and an m/z ranging from 300 to 1600. dd-MS2 scan was recorded at the 30,000 resolutions with an AGC target of 1e5, maximum injection time 100 ms, isolation window 1.4 m/z, normalized collision energy 27 and the dynamic exclusion of 30 sec.

Raw files were analysed with MaxQuant software version 1.6.0.16 for protein identification and quantification (Cox and Mann 2008). The annotation of proteins was obtained using Arabidopsis thaliana TAIR10 protein sequences (www.arabidopsis.org) plus green fluorescent protein (GFP) sequence by the integrated search again Andromeda (Cox et al. 2011). The common contaminant database (cRAP, https://www.thegpm.org) was also included in the search. The settings used for a search: 10 ppm peptide mass tolerance; 0.8 Da MS/MS tolerance; maximum of two missed cleavages allowed threshold for validation of peptides set to 0.01 using a decoy database, carbamidomethylation of cysteine was set as a fixed modification and oxidation of methionine was set as variable modification. Furthermore, the minimum peptide length of six amino was set and at least two unique peptides were required per protein group. The “label-free quantification (LFQ)” was applied to the data. Contaminants and decoy hits were removed from a dataset. The fold change (FC) was calculated by dividing the mean abundance of proteins (LFQ values in the three biological replicates) for each dehydrin LEA protein bait sample by the mean abundance of proteins (LFQ values in the three biological replicates) for the free GFP negative control sample (*pLEA10::GFP*). Only those with a FC over 1.5 were considered for further analyses. Gene Ontology analyses of interactors were performed in R with *org.At.tair.db v3.16.0* and *ClusterProfiler* packages, using with the complete Arabidopsis genome annotation as universe (background).

### Split-LUCIFERASE complementation assay

cDNA sequences of the bait and pray proteins were cloned into P221 vectors (Gateway), which were recombined into its corresponding vector for Split-Luciferase assays. List of constructs: BH-lea5-Cnluc, ZKlea4-Nnluc, ZKlea5-Nnluc, ZKosca3-Nnluc, ZKost2-Nnluc, ZKpld*α*1-Nnluc, ZKpip2b-Nnluc, ZKgr1-Nnluc, ZKlpt1-Nnluc, ZKlea8-Nnluc, ZKlea44-Nnluc, ZKlea11-Nnluc. The constructs were transformed into plant cell culture with both Kanamycin and Hygromycin selection. For luminescence detection, cell cultures were diluted to OD_600_=1. The 90 μl of transformed cell culture and PSBD was transformed into 96-well white plate and added 10 μl of diluted Nano luciferase substrate (N157A, Promega, Madison, WI) to each well. The plate was kept for 4 min at room temperature in the dark before measuring the luminescence using a CLARIOstar microplate reader (BMG LABTECH, Ortenberg, Germany) with 10-s integration periods at emission 460±30 nm with four technical replicates. The interaction pair of PFK/FBA8 as a positive control, while the pair of FBP/FBA8 used as a negative control (Zhang et al. 2023).

### BiFC assays

All constructs were generated using Gateway-compatible destination vectors. Briefly, entry clones for transient tobacco transformation were generated by PCR amplifying the open reading frames of the LEA5, LEA10, LEA4, OSCA3, PIP2B, AHA1, and PLD*α*1 genes using gene-specific primers containing Gateway recombination sites for attB3/B2 or attB1/B4 borders. Amplified attB-products were subsequently BP-cloned into pDONR221-P1P4 for LEA5 (bait protein) and pDONR221-P3P2 for all the other genes (Thermo Fisher Scientific, Waltham, Massachusetts, US) using BP Clonase according to manufacturer protocol (Thermo Fisher Scientific, Waltham, Massachusetts, US). Primers used are described in Supplementary Table 9. The resulting entry constructs were sub-cloned into pBiFCt-2in1-CC (Grefen and Blatt 2012) by LR reactions using LR Clonase II plus (Thermo Fisher Scientific, Waltham, Massachusetts, US). The identity of all constructs was confirmed by sequencing using HA-tag Rv and Myc-tag Rv primers. To evaluate basal YFP auto-reconstitution, we fused LEA5 to the nYFP moiety (attL1/4 position) but not cYFP (attL3/2 position). *Nicotiana benthamiana* plants (Tobacco) were grown for five weeks in fully controlled growth chambers with constant light intensity and long-day photoperiod (16 h day length, with a light intensity of 250 µm m−2 s−1, 22°/18°C Day/night) until transformation. Transient tobacco transformation was carried out using *Agrobacterium tumefaciens* GV3101 strain harbouring the pBiFCt-2in1-CC expression vectors described and pBintra61:p19 plasmid (Baulcombe laboratory at the plant Science Department of Cambridge University). Briefly, the *Agrobacterium tumefaciens* GV3101 strains were grown in Luria Broth at 28 °C with constant shaking at 200 rpm overnight to an OD600 of 2-4. Cells were collected by centrifugation at 2000 g, resuspended in 3 mL infiltration buffer (10 mM MES–KOH (pH 5.7), 10 mM MgCl2, 200 μM acetosyringone (3′,5′-dimethoxy-4′-hydroxyacetophenone) and incubated for 4 h at room temperature under dark conditions and gentle shaking. Induced Agrobacterium cells carrying the constructs were co-infiltrated with cells from an *Agrobacterium tumefaciens* (GV3101) strain harboring the p19 suppressor of gene silencing (Garabagi et al. 2012) in a 1:1 ratio in the abaxial side of five weeks old *N. benthamiana* leaves. Plants were kept dark at 22-23 °C for three days before confocal analysis. Confocal laser scanning microscopy (CLSM) and image processing were carried as described below. Protein-protein interaction analyses were performed in transiently transformed tobacco leaves using the ratiometric system between the fluorescence of YFP and RFP as described in (Grefen and Blatt 2012). Confocal images were acquired in a Leica TCS SP5 microscope with a 20x objective. YFP, RFP and chlorophyll fluorescence was detected using excitation/emission wavelengths of 512/520-540 nm, 566/600 nm and 488/680-720 nm, respectively. All images were acquired using sequential scanning settings. For rBiFC, YFP and RFP fluorescence signals were harvested using single planes. Imaging conditions were kept identical for all samples and experiments. The image processing was performed using Image J (FIJI) software (Schindelin et al. 2012) including area, integrated density, and limit to threshold parameters. The reconstituted YFP fluorescence is shown as proportional to the RFP cytoplasmic signal (YFP/RFP). Each experiment was repeated three times under identical conditions. Data were plotted as means ± SEM of 20 images randomly selected over the surface of four-leaf samples.

## SUPPLEMENTARY FIGURES

**Supplementary Figure 1. *LEA4*, *LEA5* and *LEA10* mRNA expression patterns in different developmental stages**

(A) Heat Map viewer displays the expression levels across 350+ samples of *LEA4*, *LEA5* and *LEA10* genes along with the corresponding subcellular localizations. (B) High-resolution spatiotemporal map of Arabidopsis primary root shows mRNA expression levels of *LEA4*, *LEA5* and *LEA10* genes in different root cell-types isolated through fluorescence-activated cell sorting (Brady et al. 2007). Root eFP browser based on the original version (Winter et al. 2007). (C) Data from Gene Expression Map of Arabidopsis show mRNA expression levels of *LEA4*, *LEA5* and *LEA10* genes in different plant developmental tissues and stages (Schmid et al. 2005). (A, B, C) Images generated and extracted from http://bar.utoronto.ca/eplant/ by (Waese et al. 2017).

**Supplementary Figure 2. *LEA4*, *LEA5* and *LEA10* mRNA expression patterns in response to salt and ABA treatment**

(A) Spatiotemporal map of five-day-old Arabidopsis primary roots shows mRNA expression levels of *LEA4*, *LEA5* and *LEA10* genes in control conditions and in response to 140 mM NaCl salt stress for 1 h (Dinneny et al. 2008). For the spatial analyses, cell type data were generated by fluorescence-activated cell sorting of roots (Dinneny et al. 2008). (B) Leaves of 5-week-old Arabidopsis plants from epidermal peels in which guard cells were the only living cell type shows mRNA expression levels of *LEA4*, *LEA5* and *LEA10* genes in control conditions and in response to 50 µm ABA for 3 h (Pandey et al. 2010). (A, B) Images generated and extracted from http://bar.utoronto.ca/eplant/ by (Waese et al. 2017).

**Supplementary Figure 3. *LEA44* and *LEA51* mRNA expression patterns in response to salt and ABA treatment**

(A) Spatiotemporal map of five-day-old Arabidopsis primary roots shows mRNA expression levels of *LEA44* and *LEA51* genes in control conditions and in response to 140 mM NaCl salt stress for 1 h (Dinneny et al. 2008). For the spatial analyses, cell type data were generated by fluorescence-activated cell sorting of roots (Dinneny et al. 2008). (B) Leaves of 5-week-old Arabidopsis plants from epidermal peels in which guard cells were the only living cell type shows mRNA expression levels of *LEA4*4 and *LEA51* genes in control conditions and in response to 50 µm ABA for 3 h (Pandey et al. 2010). (A, B) Images generated and extracted from http://bar.utoronto.ca/eplant/ by (Waese et al. 2017).

**Supplementary Figure 4. Protein levels of the dehydrins LEA44 and LEA51 and the non-dehydrins LEA1 and LEA27 in response to salt and ABA treatment**

Immunoblot of total protein extracts from (A) *pLEA1:LEA1-GFP,* (B) *pLEA44:LEA44-GFP*, (C) *pLEA51:LEA51-GFP* and (D) *pLEA27:LEA27-GFP* 10-day-old Arabidopsis seedlings in response to salt stress (150 mM NaCl) or ABA treatment (10 µM). Samples were collected at different time points of the treatment as indicated. For the western blot, anti-GFP antibodies were used in total protein extracts of T3 homozygous translational lines. Top row: immunoblot with antiGFP primary antibody. Bottom row: Ponceau S staining of the Rubisco large subunit (rbcL) as a control of protein loading and transfer. For the western blot, anti-GFP antibodies were used in total protein extracts of T3 homozygous. LEA1 and LEA27 do not belong to the dehydrin family protein. LEA1 (AT1G01470) molecular weight is 16,54 KDa, LEA44 (AT4G38410) molecular weight is 18,25 KDa, LEA51 (AT5G66400) molecular weight is 18,46 KDa, LEA27 (AT2G46140) molecular weight is 17,85 KDa and GFP molecular weight is 25 KDa.

**Supplementary Figure 5. Characterization of T-DNA insertional lines for loss-of-function mutants analysed in this study**

Expression analysis of the different candidate genes by RT qPCR of the loss-of-function mutants relative expression levels (UBQ30). Two independent alleles have been screened. 10-day-old seedlings grown in MS 0.5. Results are average from three independent biological replicates. (*p-value<0.05; **p-value<0.01).

**Supplementary Figure 6. Germination rates and primary root length of *lea5* and other loss-of-function mutants in control conditions**

(A) Germination rates of Arabidopsis seeds after four days of germination in control conditions. Data from four independent biological replicates were analysed. At least 50 seeds were quantified per genotype, and condition in each biological replicate. Shapiro-Wilk normality test revealed the normality of the data for each genotype and condition. No differences were found between genotypes in control conditions. Different letters represent significant differences in a two-way ANOVA plus Tukey’s HSD test (*p*-value < 0.05). (B) Root length of Arabidopsis seedlings after eight days of growing in control conditions. Data from four independent biological replicates were analysed. At least 20 roots were measured per genotype, and condition in each biological replicate. Boxplots depict the root length in control conditions. Shapiro-Wilk normality test revealed the normality of the data for each genotype and condition. No differences were found between root length in control conditions. Different letters represent significant differences in a one-way ANOVA plus Tukey’s HSD test with a *p*-value < 0.05).

## SUPPLEMENTARY TABLES

**Supplementary Table 1.** LEA nomenclature which enumerates genes successively on the genome (LEA1-LEA51), AGI code of the protein, alternative names of the proteins and predicted localization for the 51 LEA members in Arabidopsis.

**Supplementary Table 2.** Total number of proteins detected within the *pLEA4:LEA4-GFP* immunoprecipitates.

**Supplementary Table 3.** Total number of proteins detected within the *pLEA5:LEA5-GFP* immunoprecipitates.

**Supplementary Table 4.** Total number of proteins detected within the *pLEA10:LEA10-GFP* immunoprecipitates.

**Supplementary Table 5.** Candidate protein interactors of LEA4 enriched in the *pLEA4:LEA4-GFP* immunoprecipitates vs the GFP negative control.

**Supplementary Table 6.** Candidate protein interactors of LEA5 enriched in the *pLEA5:LEA5-GFP* immunoprecipitates vs the GFP negative control.

**Supplementary Table 7.** Candidate protein interactors of LEA10 enriched in the *pLEA10:LEA10-GFP* immunoprecipitates vs the GFP negative control.

**Supplementary Table 8.** High confident LEA5 protein interactors selected for further analyses.

**Supplementary Table 9.** Complete list of primers used for cloning, sequencing, genotyping and gene expression analysis used in this study.

## ACKNOWLEDGEMENTS

The authors thank Alisdair R. Fernie for continuous support and for providing the resources that made this research possible. We thank Youjun Zhang for performing the split-luciferase assays. N.F. acknowledges financial support from Ramon y Cajal research grant (RYC2022-038173-I) funded by Ministerio de Ciencia, Innovación y Universidades (MCIN/AEI/10.13039/501100011033) and Fondo Social Europeo Plus(FSE+).

