## Supplementary Figures for "Identification of dehydrin protein complexes *in vivo* reveals functional interactions of LEA5 with OSCA3, PIP2B and PLD*α*1 in plant water-deficit stress"

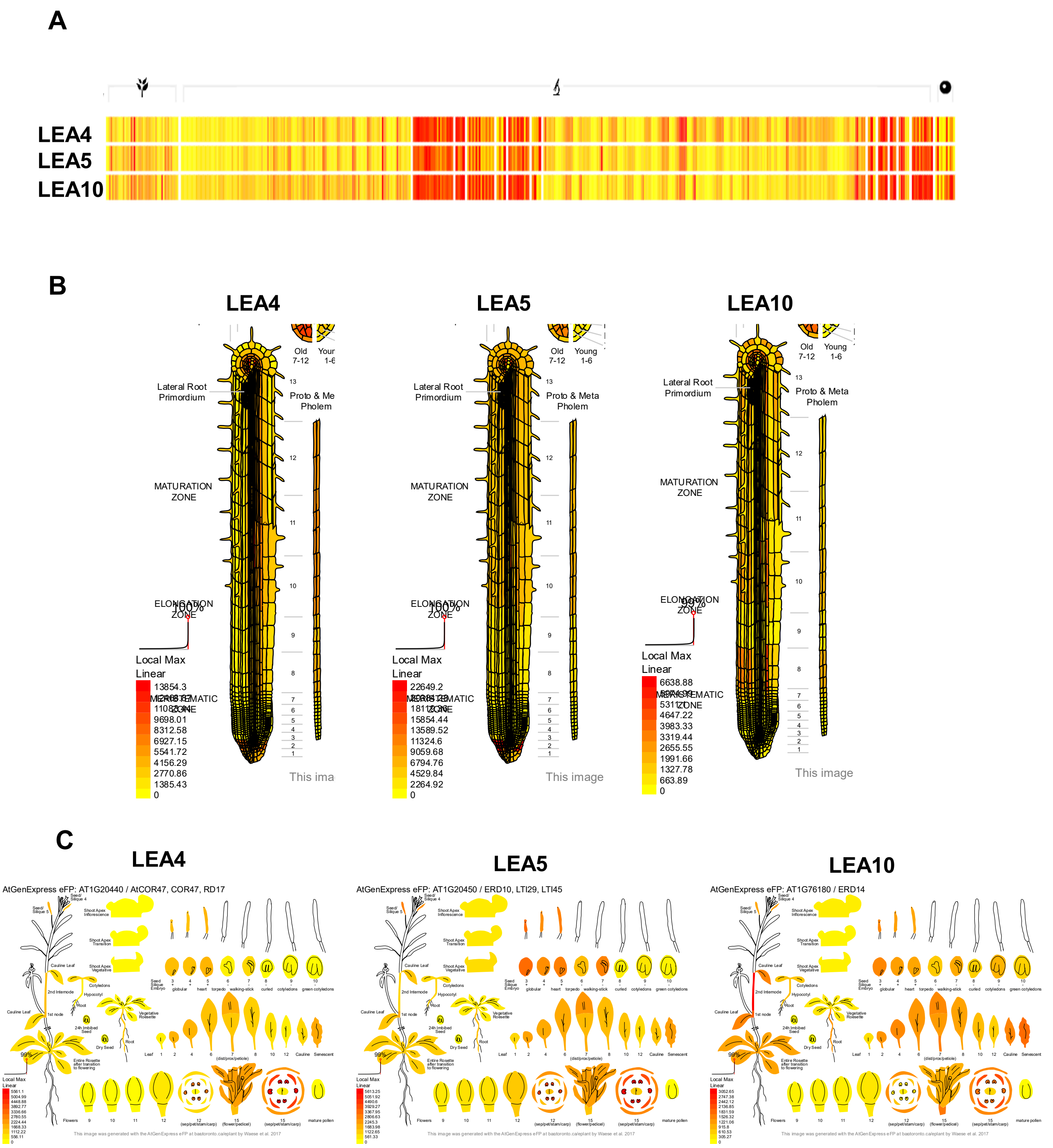


**Supplementary Figure 1. *LEA4*, *LEA5* and *LEA10* mRNA expression patterns in different developmental stages**

(A) Heat Map viewer displays the expression levels across 350+ samples of *LEA4*, *LEA5* and *LEA10* genes along with the corresponding subcellular localizations. (B) High-resolution spatiotemporal map of Arabidopsis primary root shows mRNA expression levels of *LEA4*, *LEA5* and *LEA10* genes in different root cell-types isolated through fluorescence-activated cell sorting (Brady et al. 2007). Root eFP browser based on the original version (Winter et al. 2007). (C) Data from Gene Expression Map of Arabidopsis show mRNA expression levels of *LEA4*, *LEA5* and *LEA10* genes in different plant developmental tissues and stages (Schmid et al. 2005). (A, B, C) Images generated and extracted from http://bar.utoronto.ca/eplant/ by (Waese et al. 2017).

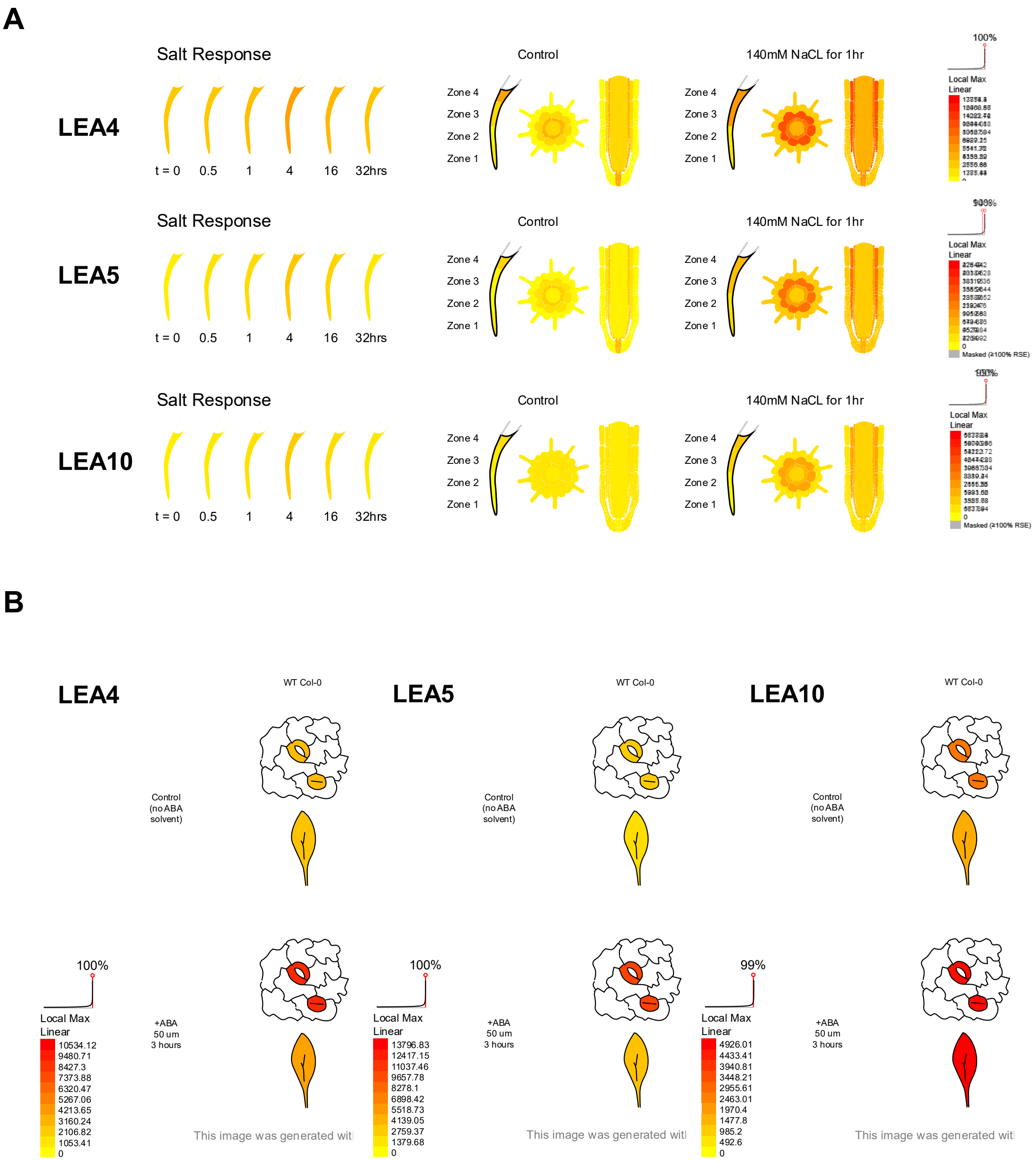


**Supplementary Figure 2. *LEA4*, *LEA5* and *LEA10* mRNA expression patterns in response to salt and ABA treatment**
(A) Spatiotemporal map of five-day-old Arabidopsis primary roots shows mRNA expression levels of *LEA4*, *LEA5* and *LEA10* genes in control conditions and in response to 140 mM NaCl salt stress for 1 h (Dinneny et al. 2008). For the spatial analyses, cell type data were generated by fluorescence-activated cell sorting of roots (Dinneny et al. 2008). (B) Leaves of 5-week-old Arabidopsis plants from epidermal peels in which guard cells were the only living cell type shows mRNA expression levels of *LEA4*, *LEA5* and *LEA10* genes in control conditions and in response to 50 µm ABA for 3 h (Pandey et al. 2010). (A, B) Images generated and extracted from http://bar.utoronto.ca/eplant/ by (Waese et al. 2017).

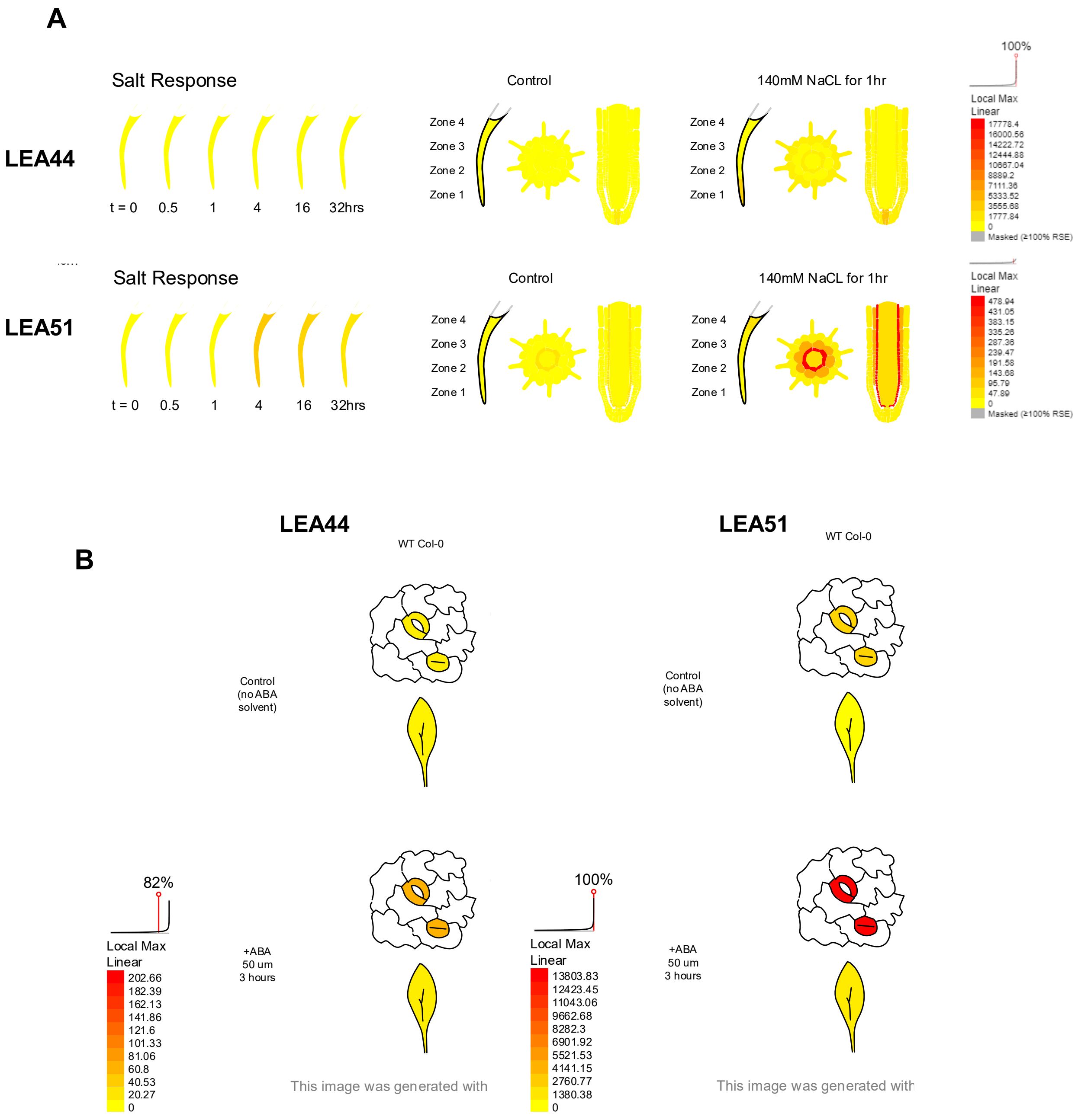


**Supplementary Figure 3. *LEA44* and *LEA51* mRNA expression patterns in response to salt and ABA treatment**

(A) Spatiotemporal map of five-day-old Arabidopsis primary roots shows mRNA expression levels of *LEA44* and *LEA51* genes in control conditions and in response to 140 mM NaCl salt stress for 1 h (Dinneny et al. 2008). For the spatial analyses, cell type data were generated by fluorescence-activated cell sorting of roots (Dinneny et al. 2008). (B) Leaves of 5-week-old Arabidopsis plants from epidermal peels in which guard cells were the only living cell type shows mRNA expression levels of *LEA44* and *LEA51* genes in control conditions and in response to 50 µm ABA for 3 h (Pandey et al. 2010). (A, B) Images generated and extracted from http://bar.utoronto.ca/eplant/ by (Waese et al. 2017).

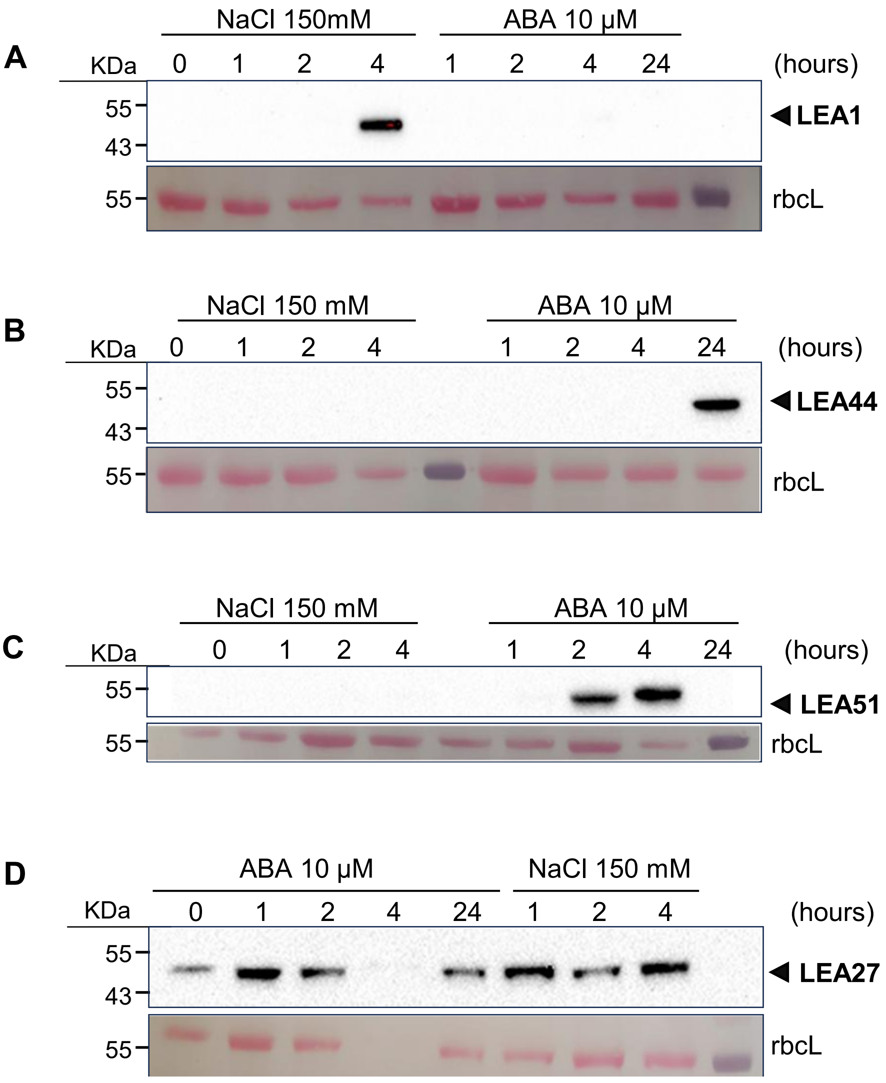


**Supplementary Figure 4. Protein levels of the dehydrins LEA44 and LEA51 and the non-dehydrins LEA1 and LEA27 in response to salt and ABA treatment**

Immunoblot of total protein extracts from (A) *pLEA1:LEA1-GFP*, (B) *pLEA44:LEA44-GFP*, (C) *pLEA51:LEA51-GFP* and (D) *pLEA27:LEA27-GFP* 10-day-old Arabidopsis seedlings in response to salt stress (150 mM NaCl) or ABA treatment (10 µM). Samples were collected at different time points of the treatment as indicated. For the western blot, anti-GFP antibodies were used in total protein extracts of T3 homozygous translational lines. Top row: immunoblot with antiGFP primary antibody. Bottom row: Ponceau S staining of the Rubisco large subunit (rbcL) as a control of protein loading and transfer. For the western blot, anti-GFP antibodies were used in total protein extracts of T3 homozygous. LEA1 and LEA27 do not belong to the dehydrin family protein. LEA1 (AT1G01470) molecular weight is 16,54 KDa, LEA44 (AT4G38410) molecular weight is 18,25 KDa, LEA51 (AT5G66400) molecular weight is 18,46 KDa, LEA27 (AT2G46140) molecular weight is 17,85 KDa and GFP molecular weight is 25 KDa.

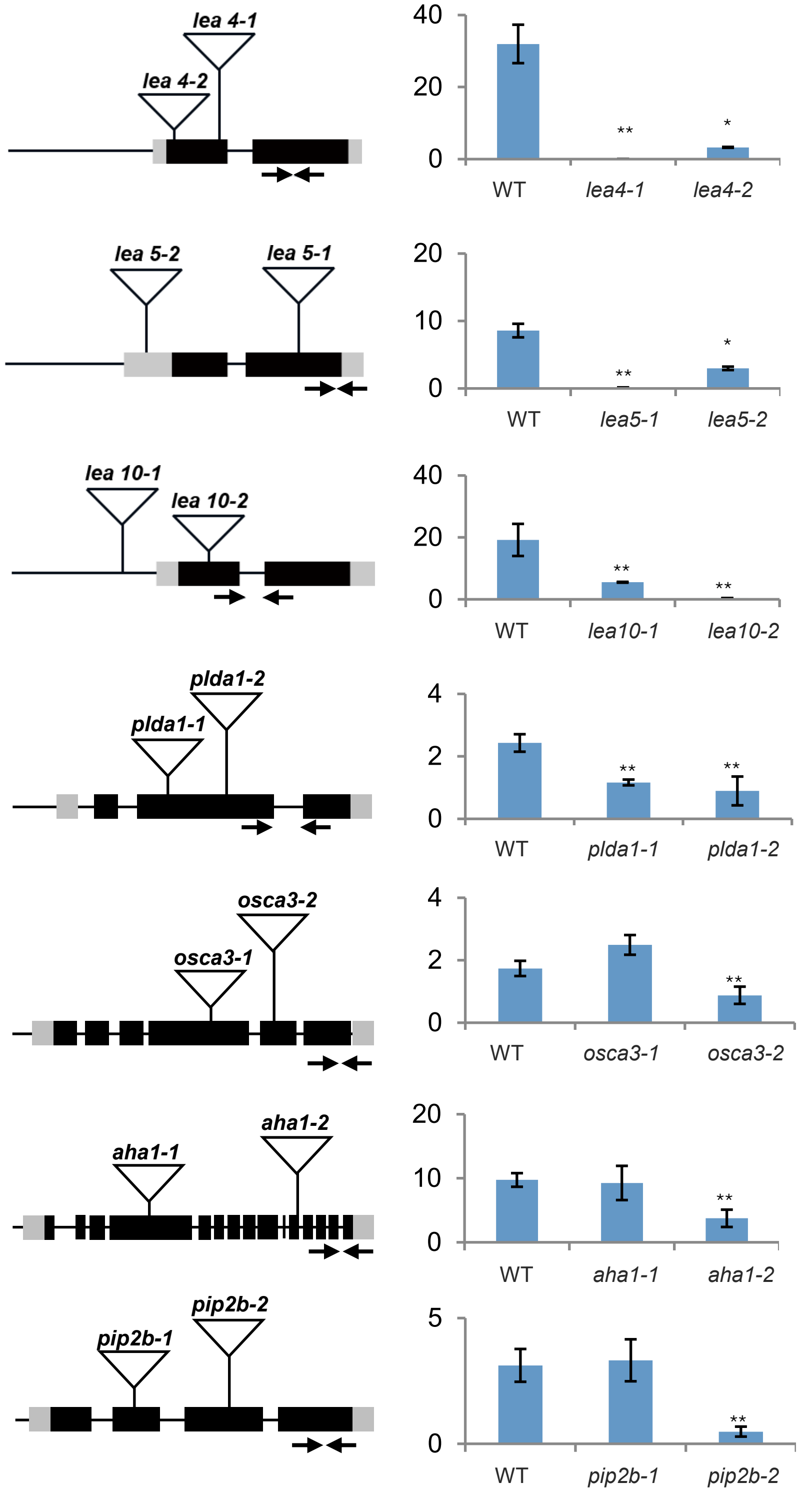


**Supplementary Figure 5. Characterization of T-DNA insertional lines for loss-of-function mutants analysed in this study**

Expression analysis of the different candidate genes by RT qPCR of the loss-of-function mutants relative expression levels (UBQ30). Two independent alleles have been screened. 10-day-old seedlings grown in MS 0.5. Results are average from three independent biological replicates. (*p-value<0.05; **p-value<0.01).

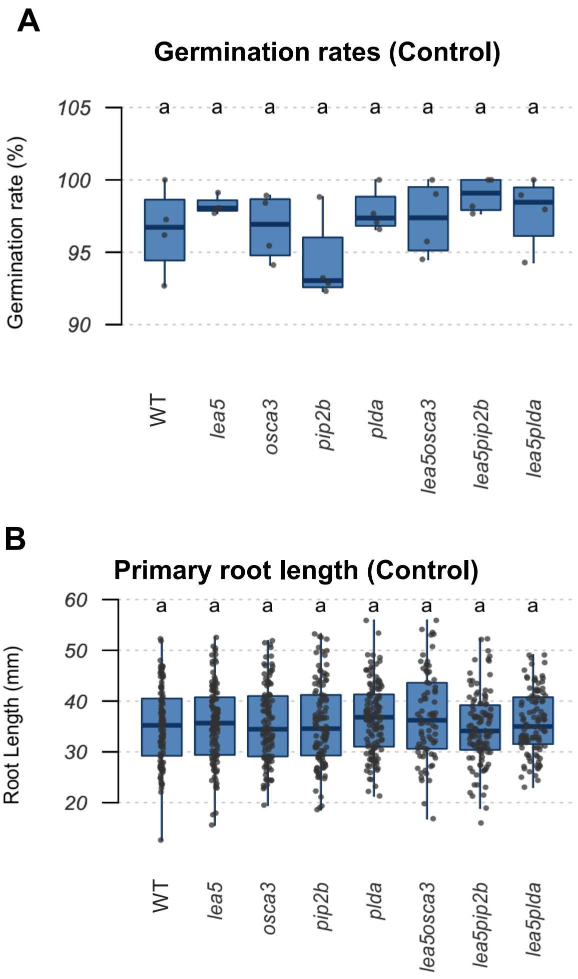


**Supplementary Figure 7. Germination rates and primary root length of *lea5* and other loss-of-function mutants in control conditions**

(A) Germination rates of Arabidopsis seeds after four days of germination in control conditions. Data from four independent biological replicates were analysed. At least 50 seeds were quantified per genotype, and condition in each biological replicate. Shapiro-Wilk normality test revealed the normality of the data for each genotype and condition. No differences were found between genotypes in control conditions. Different letters represent significant differences in a two-way ANOVA plus Tukey's HSD test (p-value < 0.05). (B) Root length of Arabidopsis seedlings after eight days of growing in control conditions. Data from four independent biological replicates were analysed. At least 20 roots were measured per genotype, and condition in each biological replicate. Boxplots depict the root length in control conditions. Shapiro-Wilk normality test revealed the normality of the data for each genotype and condition. No differences were found between root length in control conditions. Different letters represent significant differences in a one-way ANOVA plus Tukey's HSD test with a p-value < 0.05 ).
